# Diverse classes of constraints enable broader applicability of a linear programming-based dynamic metabolic modeling framework

**DOI:** 10.1101/2021.05.13.444091

**Authors:** Justin Y. Lee, Mark P. Styczynski

**Affiliations:** School of Chemical & Biomolecular Engineering, Georgia Institute of Technology, Atlanta, GA, USA

## Abstract

Current metabolic modeling tools suffer from a variety of limitations, from scalability to simplifying assumptions, that preclude their use in many applications. We recently created a modeling framework, LK-DFBA, that addresses a key gap: capturing metabolite dynamics and regulation while retaining a potentially scalable linear programming structure. Key to this framework’s success are the linear kinetics and regulatory constraints imposed on the system. However, while the linearity of these constraints reduces computational complexity, it may not accurately capture the behavior of many biochemical systems. Here, we developed three new classes of LK-DFBA constraints to better model interactions between metabolites and the reactions they regulate. We tested these new approaches on several synthetic and biological systems, and also performed the first-ever comparison of LK-DFBA predictions to experimental data. We found that no single constraint approach was optimal across all systems examined, and systems with the same topological structure but different parameters were often best modeled by different types of constraints. However, we did find that the optimal constraint approach was generally robust to local perturbations of the system, indicating that just a single wild-type dataset could allow identification of the ideal constraint for a given system. These results suggest that the availability of multiple constraint approaches will allow LK-DFBA to model a wider range of metabolic systems.

## Introduction

Mathematical and computational models are often used to study metabolism, the set of reactions that supply the chemical precursors necessary for almost all cellular processes. These metabolic models are significantly cheaper and faster to run than laboratory experiments, meaning that they can be of tremendous value when they are able to predict how changes in or to a metabolic system can affect its state. While a few pathways and sections of metabolism (e.g., glycolysis and central carbon metabolism) have been modeled and characterized quite well in a few organisms (e.g., *Saccharomyces cerevisiae* and *Escherichia coli*) [1, 2], genome-scale models that capture metabolism at a systems scale have been more difficult to develop. Metabolism involves many interconnected reactions and pathways, making it critical to include as much of metabolism as possible in metabolic models to better represent the system and generate accurate predictions. Metabolomics, which is the systems-scale measurement of metabolites in biological systems, thus has great potential to provide the information necessary to drive systems-scale metabolic models since metabolomics data are easier to acquire than, for example, metabolic flux data. However, creating genome-scale metabolic models that capture critical system behaviors like metabolic dynamics remains an outstanding challenge in the field, which has prevented the value of metabolomics data in this context from being fully realized.

The most popular type of frameworks for metabolic modeling are constraint-based models, including flux balance analysis (FBA) [3, 4], and ordinary differential equation (ODE) models. FBA assumes that the metabolic system is at steady state, which allows it to be modeled as a linear program (LP) that can be efficiently solved (even at the genome scale) but precludes modeling metabolite dynamics without substantial changes to the framework. ODE models are more widely used when dynamics are important, but are typically limited to smaller-scale modeling of well-studied pathways (e.g. central carbon metabolism [1] or glycolysis [5]) and the best-studied organisms (e.g. CHO cells [6]) due to uncertainty in the mathematical form and parameter values for the reaction kinetics terms. Only a few exceptions [7-10] have approached genome-scale ODE models, and they still require lengthy parameter estimation steps, prior information about kinetic constants, or have only been shown to be useful near the reference state of the system. As a result, steady-state fluxes continue to be the almost exclusive focus of study for genome-scale models. Modeling frameworks that can predict various metabolic phenotypes at the genome scale in a computationally tractable way have great potential for understanding, predicting, and controlling metabolism.

To address this problem, we recently developed Linear Kinetics-Dynamic Flux Balance Analysis (LK-DFBA), a modeling strategy to efficiently track metabolite dynamics [11]. LK-DFBA combines advantages of both constraint-based and ODE models, unrolling the temporal aspect of the system into a larger stoichiometric matrix that captures metabolite dynamics while retaining a LP structure. The most critical element to accomplishing this goal is the addition of linear kinetics constraints that model the interactions between metabolites and the reactions whose fluxes they affect, including mass action kinetics and allosteric regulatory interactions. The number of parameters in LK-DFBA that need to be estimated can be fewer or easier to estimate than in ODE models due to these linear kinetics constraints. This enables LK-DFBA to potentially be applied to metabolic systems of all sizes, with a smaller increase in computational burden compared to ODE models. Furthermore, because LK-DFBA retains a linear structure, it can potentially be used with many existing metabolic modeling tools that require constraint-based models, such as OptKnock [12]. We have previously shown that LK-DFBA can outperform ODE-based modeling approaches when used in conditions most relevant to metabolomics data (low sampling frequency and high noise) [11]. A framework such as LK-DFBA that can model systems at the genome scale is essential to take full advantage of metabolomics data.

In our initial description of LK-DFBA, we explored two different approaches for model parameterization. The first approach, LK-DFBA (LR), parameterizes constraints solely via linear regression of interacting metabolite concentration and flux data. The second approach, LK-DFBA (LR+), uses the parameters from the linear regressions as initial seeding values for a secondary optimization to identify the optimal constraints for each interaction. While LK-DFBA (LR+) yields better fits to training data than LK-DFBA (LR), the latter approach estimates its parameters with trivial computational effort while still producing results that are similar in error to ODE models. As a result, LK-DFBA (LR) may be the preferable approach for the efficient construction and parameterization of metabolic models at the genome scale.

However, the overall LK-DFBA framework still has some limitations in terms of how accurately it represents the underlying biology and biochemistry of the system. For example, the linear kinetics constraints used in LK-DFBA (LR) may be viewed as crude approximations of the interactions between metabolites and fluxes, which are typically non-linear in nature. While kinetic equations found in ODE models (such as Michaelis-Menten or biochemical system theory (BST) representations [13, 14]) can capture the non-linearity of these interactions, the current linear framework in LK-DFBA cannot. Additionally, when allosteric regulatory information is considered (which LK-DFBA includes in its framework), reaction fluxes are often controlled by multiple metabolites. Currently, LK-DFBA creates separate constraints for each metabolite that controls a flux, which precludes modeling how multiple metabolites simultaneously interact with a reaction flux.

Since the linear kinetics constraints are so critical in LK-DFBA’s functioning, it is likely that improving those constraints could have a substantial impact on LK-DFBA’s ability to capture and predict biological phenomena. Accordingly, we devised three new types of kinetics constraints for LK-DFBA to account for biologically important features like non-linearity and simultaneous regulation by multiple metabolites. These new approaches were compared to the original LK-DFBA (LR) constraints by testing on synthetic model systems as well as models based on *Lactococcus lactis* and *Escherichia coli* [1, 2]. We found that the ideal constraint for a given model depended not only on its structure, but also on its set of parameter values even for the same structure. The new types of constraints allowed improved fitting of data in many tested models. We then probed these constraint approaches for their robustness to model perturbation and their ability to predict phenomena not captured in training data. We found that while different model topologies and parameter sets have different optimal constraint approaches, the same constraint approach was typically optimal across perturbations for any given model, indicating that just one set of data is sufficient for parameterization robust enough for predictions of model perturbations. We also showed that the LK-DFBA approach chosen for the *L. lactis* and *E. coli* models can be used to predict changes in several critical metabolites and fluxes in qualitative agreement with literature experimental results. The new constraint strategy and its robustness to model perturbations have promise for moving the LK-DFBA framework towards modeling increasingly large metabolic systems in the future.

## Methods

### LK-DFBA

Linear Kinetics-Dynamics Flux Balance Analysis (LK-DFBA) is a recently developed modeling strategy that is both scalable and capable of capturing metabolite dynamics. The full details of this approach have been described in detail previously [11], so we only outline the most important aspects of our framework here. In brief, LK-DFBA is a linearly-constrained quadratic program with temporal dynamics modeled by discretizing time and unrolling the system into a larger matrix *A* representing each time point separately. *A* consists of the stoichiometric matrix of the system and an identity matrix, which when multiplied by 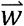 (a vector of fluxes 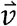 and pooling fluxes 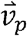 (i.e. 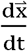)), calculate the system’s mass balances at each time point *t*_*k*_. The fluxes are constrained by lower 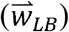 and upper 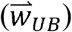 bounds. LK-DFBA includes a quadratic objective function *z* where λ is a small penalty on the norm of the solution Vector 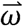 and 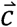 is a vector of weights. While LK-DFBA is technically a quadratic program, we found that adding this penalty reduced the possibility of solution degeneracy and led to no appreciable increase in computation time compared to a linear objective function. The solution vector consists of fluxes 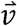 and metabolite concentrations 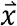 at each timepoint in the unrolled model. Like the fluxes, the metabolite concentrations are also constrained by lower 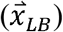 and upper 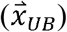 bounds. Linear inequality constraints that model mass action kinetics and metabolite-dependent regulation are included in the model; they are the driving force behind metabolite accumulation and depletion by limiting the maximum flux *v*_*i*_ allowed based on the availability of metabolites *x*_*j*_ over time. In the LK-DFBA (LR) approach, these linear kinetics constraints are modeled using linear regression on assumed available metabolomics and fluxomics data to estimate *a* and *b* parameters for each of the *n* pairings of metabolites that participate in a flux reaction. If fluxomics data are unavailable, dynamic flux estimation (DFE) can be used to infer flux values from concentration data [15]. The formulation of an LK-DFBA model is presented in Equation 1.

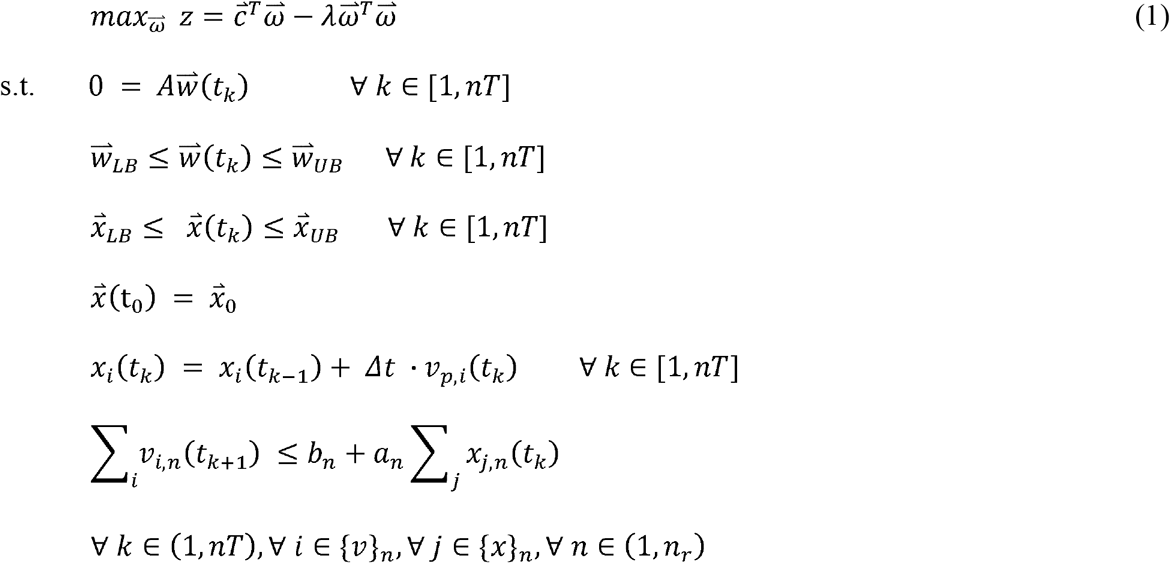

where

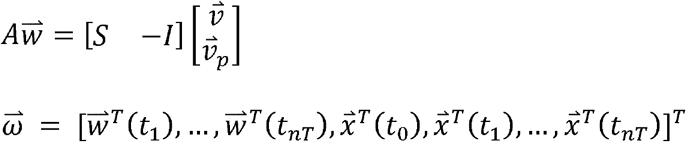

In the LK-DFBA (LR+) method, the parameters from the LK-DFBA (LR) approach are used as initial estimates in a secondary optimization step that finds improved kinetics constraint parameters, though at the cost of computational time. Because LK-DFBA retains a simple QP structure with entirely linear constraints, it is readily scalable and has the potential to be used with current constraint-based modeling tools. The objective functions used for each individual model are described in the Supplementary Methods.

### Constraint Approaches

#### LK-DFBA (LR)

The original LK-DFBA approach uses linear kinetics constraints to model the interaction between a metabolite and a flux, parameterized using linear regression on available metabolomics and fluxomics data. These constraints take the form of *v* ≤ *ax + b*, where *v* is the flux being constrained, *x* is the concentration of a metabolite that interacts with the flux, and *a* and *b* are the linear constraint parameters. These interactions may be due to mass action kinetics, where the interactions are known based on the stoichiometric topology of the system, or they may stem from allosteric regulation. While we have previously shown that these linear approximations of metabolic interactions can be effective for modeling metabolism, they are still approximations of the true non-linear and interconnected biochemical relationships in metabolism. Below, we discuss three new constraint approaches to address these potential limitations.

#### LK-DFBA (NLR)

While the key advantage of using constraint-based models is their LP structure that enables efficient identification of the optimal solution of the problem, most metabolite-flux interactions exhibit non-linear behavior that may not be captured well by linear equations. Recently, computational solvers have improved such that quadratically constrained programs (QCPs) are not much more computationally expensive than LPs. Accordingly, we implemented quadratic constraints into the LK-DFBA framework to explore their potential for improving model accuracy with only a modest increase in computational time. One important aspect of LPs and QCPs is that all of the constraints must create a convex feasible solution space to guarantee that a global optimum can be found [16]. If *v* ≤ *ax*^2^+ *bx* + *c* represents a quadratic constraint, where *v* is the flux being constrained, *x* is the concentration of a metabolite that interacts with *v*, and *a, b*, and *c* are the parameters of the quadratic constraint, *a* must be a negative value to retain a convex solution space. If *a* is found to be a positive value during parameterization, we convert the quadratic constraint into its original linear form as found in LK-DFBA (LR). The parameters were estimated using linear least squares regression. We refer to this overall approach as LK-DFBA (NLR).

#### LK-DFBA (DR)

Enzymatic reactions are often controlled by more than a single metabolite that can either induce or inhibit enzyme activity, which should ideally be captured in the model constraints. To model such regulation of a reaction by multiple metabolites, LK-DFBA (LR) creates individual linear constraints for each controller metabolite that are independent of each other and are thus unable to capture the synergistic or antagonistic effects of multiple metabolites working in conjunction to regulate a flux. We implemented a new strategy that uses dimensionality reduction to consolidate information content from all controller metabolites for a flux into a single constraint. Dimensionality reduction is often used in data analysis, including analysis of metabolomics data, to more easily represent and digest datasets with many measured variables. Principal component analysis (PCA) is one of the most commonly used dimensionality reduction approaches; it linearly transforms the original variables into new, orthogonal composite variables called principal components that capture as much variance in the original variable data in as few principal components as possible [17]. Ideally, the first or first few principal components capture the majority of the variance in the original dataset, which allows one to analyze only those composite variables rather than all of the original variables at once. Here, we use PCA to capture the maximal variance of the controller metabolite data in a single principal component and use that composite variable as the regressor during linear regression with the target flux data. These new constraints are represented as *v* ≤ *aPC*_1_ + *b*, where *v* is the flux being constrained, *PC*_*1*_ is the controller metabolite concentration data projected into the first principal component, and *a* and *b* are the constraint parameters. The parameters were estimated using linear least squares regression. We refer to this dimensionality reduction approach as LK-DFBA (DR).

#### LK-DFBA (HP)

Another approach we used to model interactions of a flux with multiple metabolites was hyperplane constraints. Unlike LK-DFBA (DR), which always builds constraints in two dimensions (i.e. the target flux vs. the first principal component), the hyperplane constraint exists in (*n* + 1) dimensions, where *n* is the number of metabolites that control a target flux. This approach may avoid loss of information content from metabolite data as is possible during dimensionality reduction: as the number of metabolites in an interaction increases, the likelihood of the first principal component not capturing the majority of variance in the data increases. The hyperplane constraint equation can be represented as 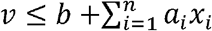, where *v* is the flux being constrained, *n* is the number of metabolites that interact with *v, x*_*i*_ is the concentration of metabolite *i, a*_*i*_ is the constraint parameter for metabolite *x*_*i*_, and *b* is another constraint parameter. The parameters were estimated using linear least squares regression. We refer to the hyperplane approach as LK-DFBA (HP).

### Test Models

#### Synthetic model

The first system we examined was a simple synthetic model with five metabolites and five fluxes that was derived from a branched pathway model used previously [11]. This system is represented via an ODE-based model that uses power-law kinetics to represent reaction fluxes [14]. The kinetic equations for each pathway are shown in Figure 1. To create a variety of synthetic models with the same stoichiometric topology, we randomly generated *a* and *b* parameters in each kinetic equation. The parameters for each model can be found in Table S1. Time course metabolite and flux data were generated by solving the ODE system in MATLAB (2018b).

**Figure 1:**
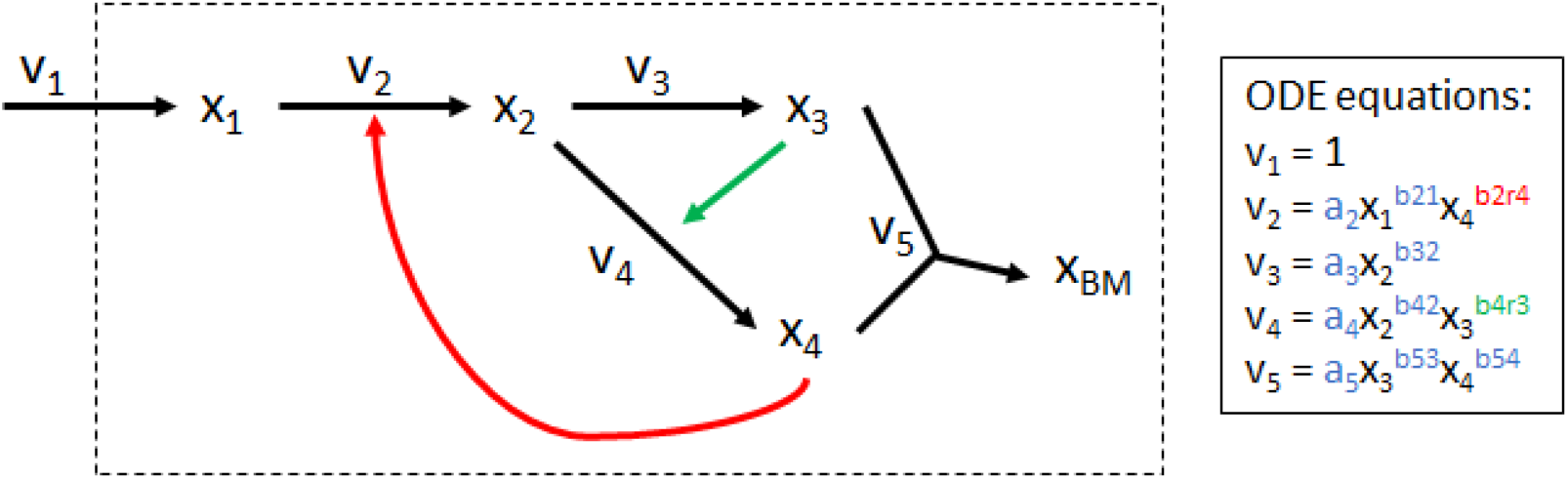
Synthetic model. Adapted from another branched pathway model used in previous work [11]. v_1_, v_2_, v_3_, v_4_, and v_5_ are system fluxes (black arrows) and x_1_, x_2_, x_3_, x_4_, and x_BM_ are metabolites, where x_BM_ is a metabolite representing biomass. Green and red arrows represent positive and negative regulatory interactions, respectively. ODE equations for the model are shown in the inset, where blue *a* and *b* parameters are mass action kinetic parameters and green and red *b* parameters are positive and negative regulatory parameters, respectively.

#### *Lactococcus lactis* model

This model was created by Costa *et al*. and comprises central metabolism and production pathways for important metabolites such as mannitol and 2,3-butanediol [2]. The *L. lactis* model consists of 26 metabolites and 21 fluxes and is publicly available on KiMoSys [18]. Noiseless data were generated in COPASI 4.24 (Build 197) using the default initial conditions and parameters over a simulation time of two hours.

#### *Escherichia coli* model

The *E. coli* model developed by Chassagnole *et al*. encompasses glycolysis and the pentose phosphate pathway [1]. This model is publicly available on KiMoSys, but was rebuilt within MATLAB to allow easy creation of new models that use the original *E. coli* model’s topology and stoichiometry. Noiseless data for the original *E. coli* model were generated in MATLAB (2018b) using the default initial conditions and parameters, while random initial conditions and parameters were used for the new models with the *E. coli* topology. To be consistent with our previous work, we used a simulation time of ten seconds [11]. More information about model parameters, the objective functions used for each model, and additional implementation details can be found in the Supplementary Methods.

### Pathway Perturbations

To test the ability of LK-DFBA to predict metabolic behaviors not represented in the training data, we introduced perturbations into each system either through down-regulation (indicated with the prefix ‘d’ in all figures) or up-regulation (indicated with the prefix ‘u’) of reaction fluxes. For the synthetic models, we down-regulated v_2_, v_3_, and v_4_ by multiplying their constraint equation parameters (which restricts their maximum allowable flux value) by 0.5x and up-regulated these pathways by doubling the constraint equation parameters. The pathways and reactions to be perturbed in the *L. lactis* [2, 21-24] and *E. coli* [25-29] models were chosen based on previous literature. Reactions in the *L. lactis* model (lactate dehydrogenase, phosphofructokinase, acetate kinase, mannitol 1-phosphatase) were down-regulated to 0.1x their original parameter values (since completely knocking out reactions would often produce infeasible solutions for the linear program) and up-regulated to 2x their original parameter values, magnitudes that were necessary to effect significant perturbations to the system’s behavior. Reactions in the *E. coli* model (pyruvate kinase, phosphoglucose isomerase, glyceraldehyde-3-phosphate dehydrogenase, phosphofructokinase, triose-phosphate isomerase, ribulose-phosphate epimerase, phosphoglucomutase) were down-regulated to 0.1x and up-regulated to 2x their original parameter values.

### Generating Noisy Data

Noise was introduced into the system using the original noiseless data (50 time points) and a down-sampling of that dataset into nT time points evenly spaced over the time interval of interest. Both metabolite and flux values were then replaced with a random value drawn from the normal distribution *N*_*i,k*_, ∼ (*y*_*i*_ (*t*_*k*_),*CoV*·*y*_*i*_ (*t*_*k*_)), where *y*_*i*_ (*t*_*k*_) is the value of species (metabolite or flux) *i* at time point *k*, and CoV is a coefficient of variance. For each sampling frequency and CoV condition, ten noisy datasets were generated.

## Results

### Fitting and predicting phenotypes in synthetic models

We generated twenty random sets of parameters and initial conditions for the kinetic equations in the synthetic model to examine the utility of the new constraint approaches across models of the same topology but different parameter sets. We produced *in silico* metabolite concentration and flux data over a time interval of ten seconds by solving the ODEs in each synthetic system. The four constraint approaches were used for parameterization of LK-DFBA models to the twenty datasets. The fitted LK-DFBA models were then simulated over the same interval using the initial conditions for each respective synthetic system to compare against the original ODE data. This process was performed on both noiseless (nT = 50, CoV = 0) and noise-added data with different sampling frequencies (nT = 50 or 15) and levels of noise (CoV = 0.05 or 0.15).

For the noiseless cases (Figure 2A), the best-fitting constraint approach on the wild-type data (dark green highlighted in cyan boxes) varied across the different models. All four approaches performed best for at least one of the models. Given that these models all have the same topology and just different parameter values, one might have expected they would be sufficiently similar that one single type of constraint would perform best across all or most models. The fact that different constraints fit different models better may be due to different qualitative behaviors being evident in the metabolic dynamics in the different models. Since all parameters were resampled in each model, they might access fundamentally different regimes of concentration or flux values and thus be better fit by different types of constraints. Similar trends were evident in noisy data (representative example in Figure 2B), where the optimal constraint approach varied across the different models (highlighted in cyan boxes in Figure 2B).

**Figure 2:**
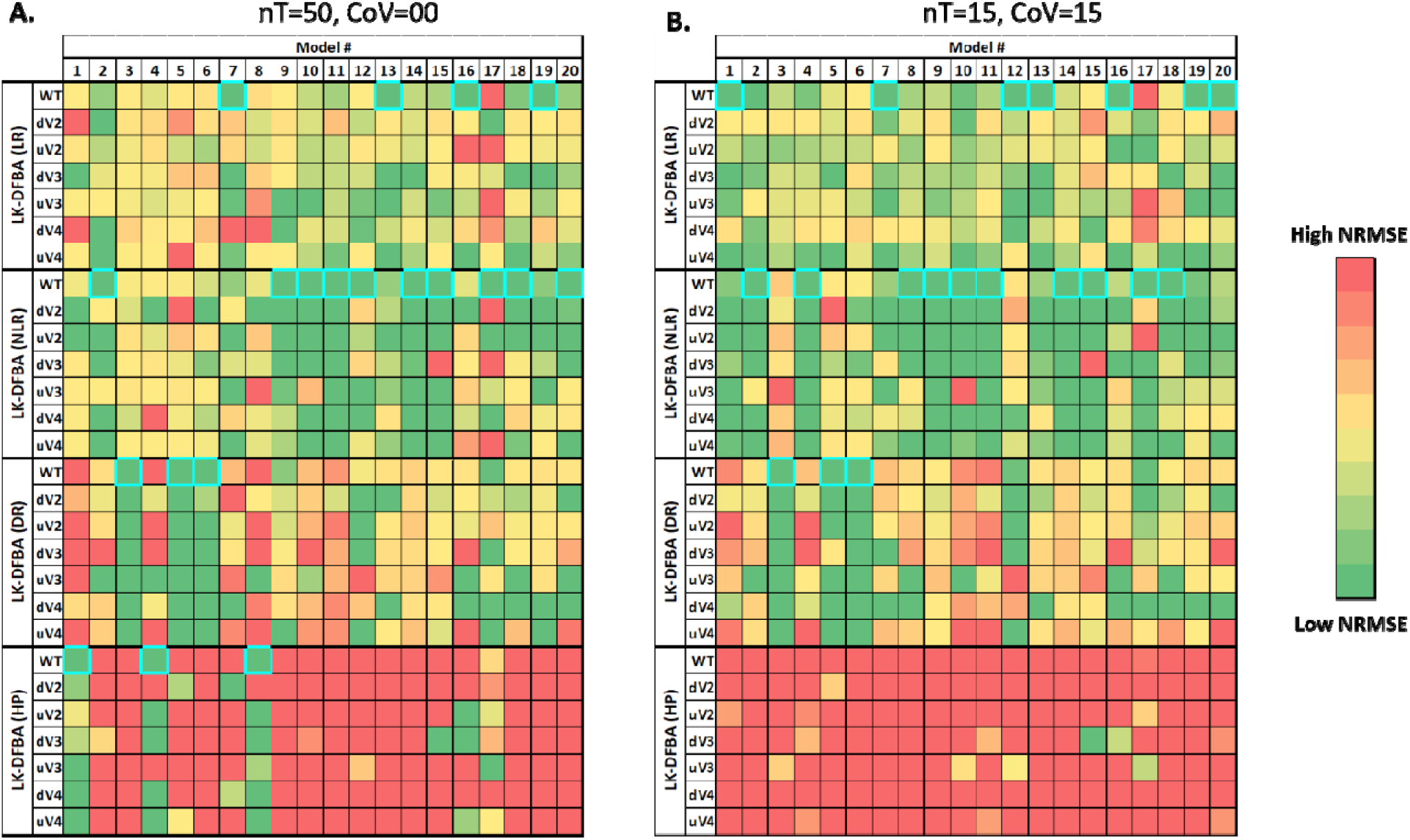
NRMSE heatmap of LK-DFBA approaches on different synthetic models. Each constraint approac was used to fit parameters to wild-type (WT) data and then used to simulate the WT system and *in silico* genetic perturbations with fluxes v_2_, v_3_, or v_4_ down- or up-regulated. The best constraint approach for each WT model is highlighted by a cyan border. Dark green boxes represent the lowest NRMSE within each genetic perturbation for each synthetic model, while dark red boxes represent the highest NRMSE (meaning that the dynamic range of the color scale varies for each perturbation for each synthetic model to better convey the relative performance of different methods). Panel A shows results for noiseless data with a sampling frequency of 50 time points. Panel B shows results for noisy datasets with a sampling frequency of 15 time points and with noise added at a CoV of 0.15. The average NRMSE of 10 noisy datasets is shown in the heatmap. The same NRMSE heatmaps with explicitly annotated error values can be found in Figure S1 and Figure S5.

Since a primary goal of building metabolic models is to predict the behavior of systems in conditions beyond those in which they are trained, we then tested the ability of each LK-DFBA model trained on wild-type data with different constraint approaches to predict the effects of defined genetic perturbations. We down- and up-regulated the v_2_, v_3_, and v_4_ pathways in the original kinetic equations by multiplying the kinetic coefficient parameters (*a* parameters in the inset of Figure 1) by 0.5x or 2x, respectively, and generating new ODE data to compare to LK-DFBA predictions. We then simulated the LK-DFBA model after adjusting the fitted LK-DFBA constraints to reflect the down- or up-regulation by multiplying the kinetics constraint parameters by 0.5x and 2x, respectively. The normalized root mean square error (NRMSE; see Supplementary Methods) between the predicted LK-DFBA metabolite concentrations and the ODE concentration data from the perturbed synthetic models was then calculated.

Using noiseless training and test data, we observed that the best constraint approach for the wild-type training data (WT) was typically also the best for the test data across the perturbations (dV2 through uV4), indicated by the clustering of dark green boxes in the rows for the constraint approach containing the highlighted cyan box for each model (Figure 2A). The NRMSE heatmap with quantitative error values can be found in Figure S1. This suggests that while global perturbations that affect the entire network (i.e., by resampling all parameter values) can often lead to different constraint approaches being optimal, local perturbations yield sufficiently similar behavior to allow the same constraint approach to be optimal for both the training and the test data.

When using noisy data, similar trends were observed (representative example in Figure 2B). While the best constraint approach for WT noisy data was not always the same as the best approach for noiseless data, the best constraint approach for a given noisy WT dataset was still generally the best for predicting the impacts of *in silico* genetic perturbations in the same model (dV2 through uV4). Interestingly, noisy data negatively affected the performance of LK-DFBA (HP) to a much greater extent than the other approaches, which caused LK-DFBA (HP) to never be identified as the best approach under the most realistic conditions of nT = 15 and CoV = 0.15 (nor for almost any other noisy data condition, except for one (Figure S2)). NRMSE heatmaps with quantitative error values for the different sampling frequencies and noise levels can be found in Figure S2-S5.

We also tested the effect of smoothing the noisy (nT = 15, CoV = 0.15) metabolite concentration and flux time course profiles by fitting to a previously described [31] impulse function (Figure S6). Smoothing the noisy data often led to lower error of the final model but required increased computation time for estimating the parameters of the impulse function and in certain cases can actually increase error if a specific dataset deviates significantly from all of the profiles that an impulse function can capture. The best constraint approach for WT smoothed data was the same as for unsmoothed data in 19 of the 20 models. As with the unsmoothed cases, the best constraint approach for smoothed data was typically consistent between WT and *in silico* genetic perturbations, and there were no cases where LK-DFBA (HP) performed the best (and it was generally the worst out of the four approaches) for smoothed data.

### Fitting and predicting phenotypes in *L. lactis* and *E. coli* models

For the *L. lactis* model, we tested the four constraint approaches on noiseless data and noisy data under different conditions (nT = 50 or 15, CoV = 0.05 or 0.15). On the noiseless data, the best constraint approach for the WT system was LK-DFBA (HP), which also had the lowest NRMSE when predicting the results of perturbations to five different genes (Figure 3A). At high sampling frequencies and low noise (nT = 50, CoV = 0.05), LK-DFBA (HP) still performed the best, but as more noise was added or lower sampling frequencies were used, LK-DFBA (NLR) overtook it to become the optimal approach. This is consistent with the findings described above for the small synthetic systems where LK-DFBA (HP) can produce low NRMSE with noiseless data but has difficulties under more realistic conditions.

**Figure 3:**
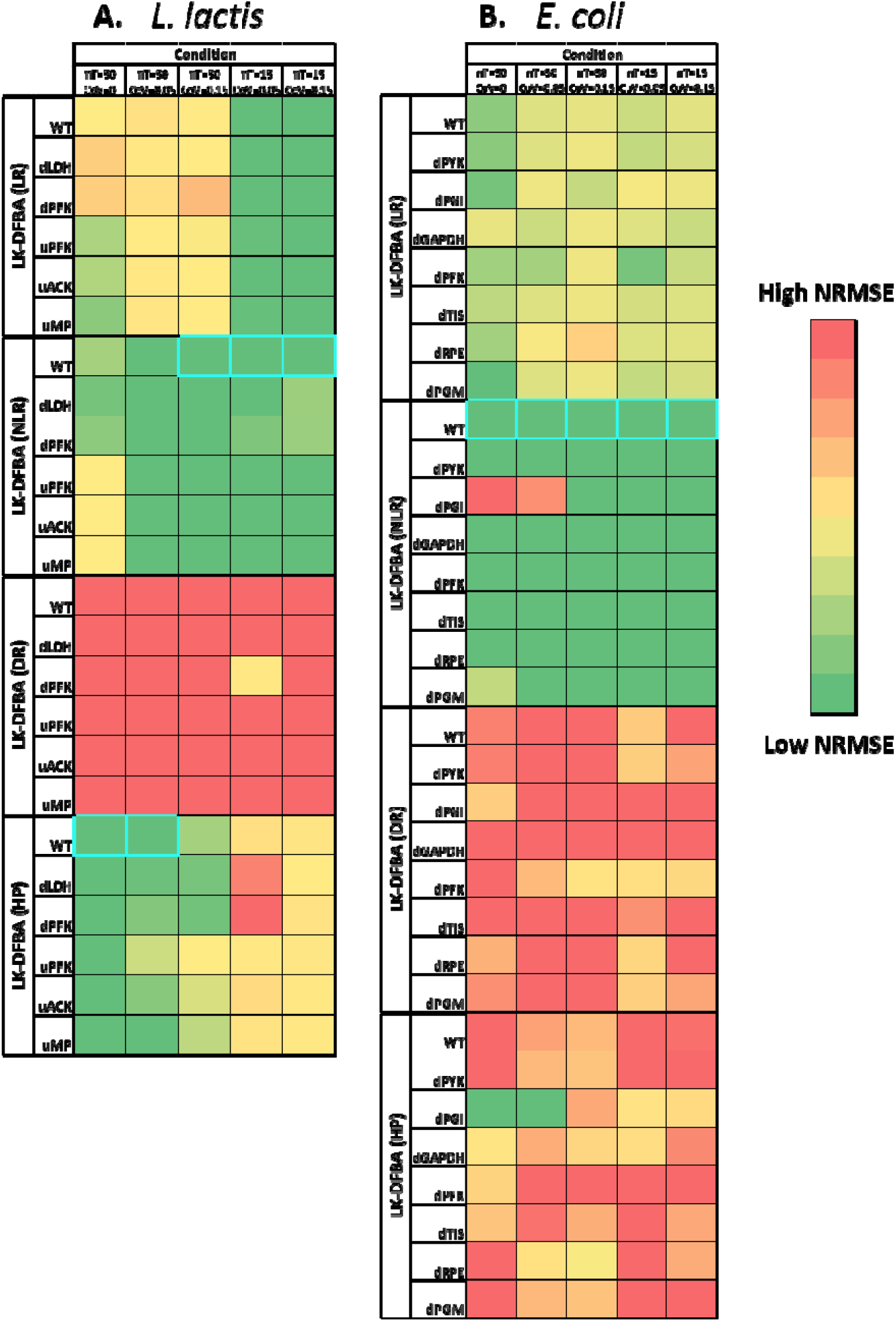
NRMSE heatmaps of constraint approaches on *L. lactis* and *E. coli* models. Each constraint approach was used to fit parameters to wild-type (WT) data and then used to simulate the WT system and the system with *in silico* genetic perturbations with literature-reported important pathways down- or up-regulated. The best constraint approach for each model is highlighted by a cyan border. Dark green boxes represent the lowest NRMSE within each phenotype for each model, while dark red boxes represent the highest NRMSE. Both the *L. lactis* (A) and *E. coli* (B) heatmaps show the mean of 10 noisy datasets, except for the noiseless condition (leftmost columns in eac heatmap). The same NRMSE heatmaps with explicitly annotated error values can be found in Figure S7.

As with the *L. lactis* model, we tested all constraint approaches on both noiseless and noisy data from the *E. coli* model under different conditions (nT = 50 or 15, CoV = 0.05 or 0.15). For this model, LK-DFBA (NLR) was the best constraint approach for noiseless data (Figure 3B). Noisy *E. coli* data produced the same results: for all noisy conditions, LK-DFBA (NLR) was optimal for the WT system. It was also optimal for almost all of the *in silico* genetic perturbations, showing once again that the same constraint approach that was optimal for the WT system at a given sampling condition was generally also optimal for the perturbed systems, supporting the generalizability of selecting a constraint approach based only on WT data.

We also perturbed the original parameters and initial conditions (drawing from the random normal distribution *N*_*i*_ ∼ (*p*_*i*_, *p*_*i*_) and taking the absolute value, where *p*_*i*_ is the original value of the *i*th parameter) of the *E. coli* model to create five new models with the same topology (more information about parameter randomization can be found in the Supplementary Methods). As with the twenty different versions of the small synthetic system, we found that across these models with identical topology and only different parameter values, the best constraint approach varied substantially (Figure S8). Again, the rates of individual reactions and how they affect overall model dynamics appear to be critical factors in determining the optimal constraint approach.

### Improved LK-DFBA predictions yield qualitative consistency with experimental *L. lactis* metabolite concentration data

To further assess how well LK-DFBA performs when predicting different phenotypes, we compared the predictions of LK-DFBA to available experimental data—the first time such a comparison has been performed for this modeling framework. The previously described ODE-based *L. lactis* model was originally parameterized using experimental metabolite time course data from *L. lactis* cultures grown with an initial glucose concentration of 40 mM [2] and validated by comparison to some experimental data from cultures grown at initial concentrations of 20mM and 80 mM glucose. Here, we similarly fitted all LK-DFBA approaches to data generated by the ODE model at 40 mM glucose and then, using the best constraint approach, simulated the LK-DFBA model at 20 mM and 80 mM initial concentrations of glucose for validation.

Figures 4A, 4B, and 4C depict the metabolite concentrations predicted by LK-DFBA (HP) (the best approach for noiseless data in the *L. lactis* model) when trained on noiseless data. For multiple initial glucose concentrations, LK-DFBA (HP) captured the general qualitative trends of glucose (depletion) and lactate (accumulation), two key metabolites in *L. lactis* that are often studied [32, 33]. For cofactor metabolites that participate in many different reactions, such as ATP, NAD(H), and inorganic phosphate (Figure S9), it was more challenging for LK-DFBA (HP) to predict their concentration profiles over the simulation interval, which is a problem found in other modeling frameworks [7]. Although LK-DFBA’s predictions were overall not as smooth or quantitatively accurate as the ODE model, this is to be expected due to the lack of a validated objective function for this constraint-based model; the objective function we used was a gross approximation that likely does not reflect the cell’s true “goal”, and it is known that the objective function can significantly affect the predictions of FBA approaches. Nevertheless, as presented here, LK-DFBA can still qualitatively track important metabolite dynamics even when using a crude objective function. This is important to note, as many organisms that are not well-studied have no readily available objective function to use.

**Figure 4:**
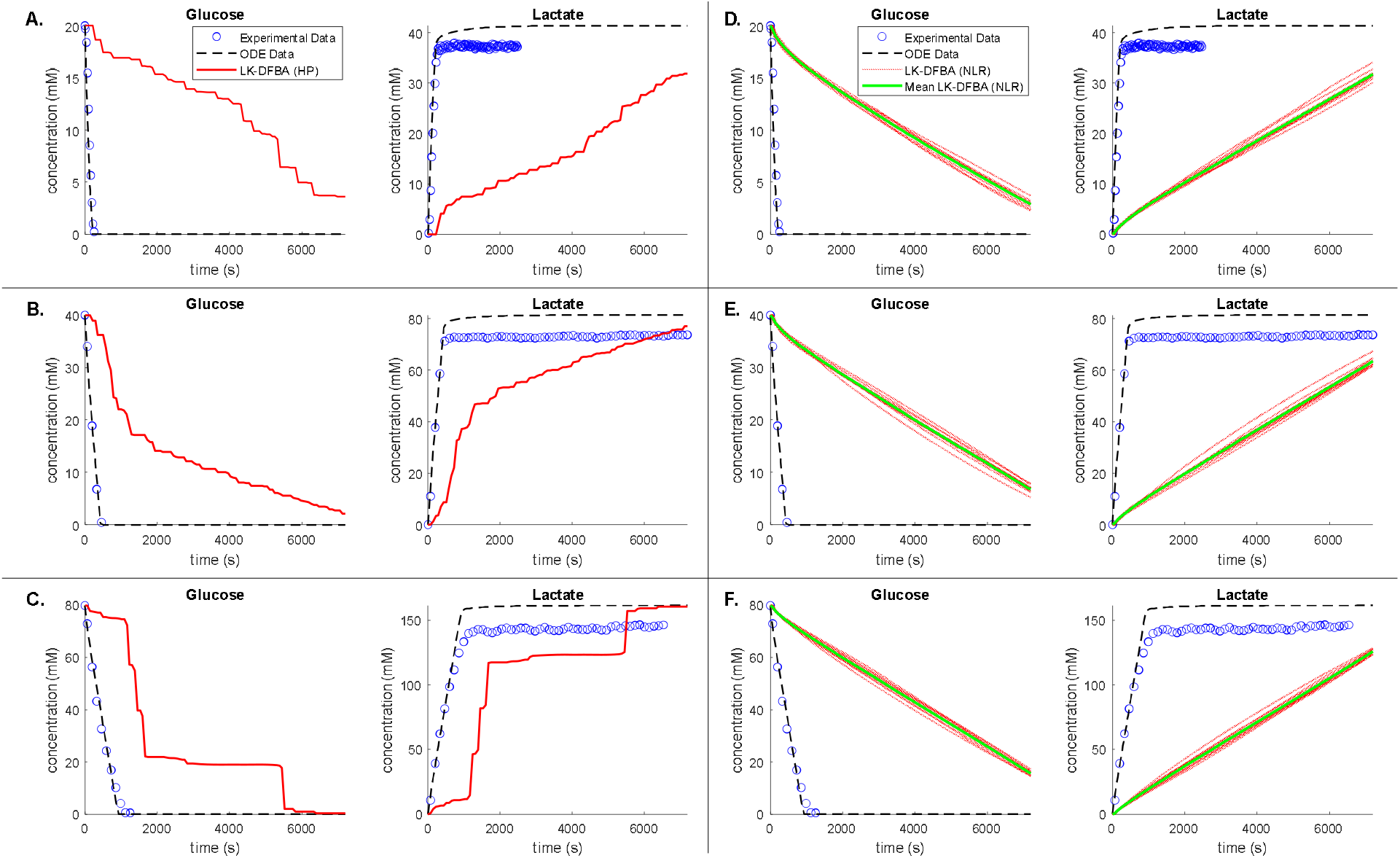
Comparison of LK-DFBA metabolite concentration predictions against ODE data and *L. lactis* experimental data when fitted to noiseless and noisy ODE data. Panels A, B, and C depict concentration profiles for LK-DFBA (HP) (solid red line) and the ODE model (dashed black line) compared to experimental data (blue circles) for initial glucose concentrations of 20 mM, 40 mM, and 80 mM, respectively, when LK-DFBA is fitted to noiseless data. Panels D, E, and F depict concentration profiles for LK-DFBA (NLR) on 10 noisy datasets (nT = 15, CoV = 0.15) and the ODE model compared to experimental data. The mean concentration profile (solid green line) is shown with each of the concentration profiles (solid red lines) from the 10 noisy datasets.

Figures 4D, 4E, and 4F depict the glucose and lactate concentration profiles predicted by LK-DFBA (NLR) (the best approach for noisy data in the *L. lactis* model) after being fitted to 10 noisy datasets generated by the ODE model and simulated at 20 mM, 40 mM, and 80 mM initial glucose, respectively. Again, the LK-DFBA framework generally captured the qualitative trends of major metabolites such as glucose and lactate, though unsurprisingly not as accurately as when noiseless data are used and with difficulties predicting cofactor concentrations (Figure S10). Because LK-DFBA (NLR) contains quadratic constraints, its results are generally smoother compared to the other LK-DFBA approaches, which helped it predict some metabolites, such as PEP, arguably better than in the noiseless case. Furthermore, LK-DFBA (NLR) is less susceptible to noise for some metabolites, such as glucose and lactate, as observed in predicting similar time courses across the 10 noisy datasets. This could be advantageous if one is modeling a system with multiple noisy data sets and requires consistent predictions for certain metabolites. Likewise, if only using a single dataset, LK-DFBA (NLR) can ensure that these metabolic profiles would not dramatically change if a different dataset had been used. Other methods, such as the original LK-DFBA (LR) approach, can result in more varied predictions (Figure S11) depending on the noisy dataset used; some appear to produce better predictions than LK-DFBA (NLR), while others are worse (though all predictions follow the same trends). These observations reiterate that the best approach is dependent on the systems and datasets being studied, so having multiple LK-DFBA approaches available is an improvement over only using the LK-DFBA (LR) framework.

### Changes in LK-DFBA flux profiles due to gene knockouts are correlated with experimental *E. coli* steady-state flux data

We also compared the predictions of the best LK-DFBA approach on the *E. coli* model to experimental steady-state flux data obtained through gene knockout experiments by Ishii *et al*. [26]. Because the Chassagnole model, which LK-DFBA is fitted to, only encompasses central carbon metabolism, we focused on 13 gene knockouts and 14 fluxes that are included in both the Chassagnole model and the Ishii steady-state flux results. We used the dilution rate of 0.2 h^-1^ for all experimental data. To emulate a gene knockout in the LK-DFBA model, we down-regulated the pathway(s) that correspond with the gene by multiplying the parameters of the relevant constraints by 0.1x instead of completely removing the reaction, as we found that this sufficiently reduced the possible flux reaction rate without causing infeasible solutions from the solver. Additionally, it is not uncommon for enzymatic activity to remain in a pathway after single gene knockouts due to paralogous enzymes and enzyme promiscuity. Because the LK-DFBA predictions do not reach steady-state for the simulation time examined in this work and our previous work (ten seconds), we instead used the average flux of the predicted time course to describe how LK-DFBA’s predictions change from the wild-type to gene knockout phenotype (see Methods). The average flux before and after a gene knockout should reflect whether the reaction rate generally increases or decreases across the studied time interval after a system perturbation. We used a Pearson correlation to determine if the average flux profiles of 14 reactions predicted by LK-DFBA changed similarly to the experimental data after a gene knockout (see Supplementary Methods for details). This assessment method has been used previously by Lima et al. to compare multiple *E. coli* models, including the Chassagnole model, to the Ishii dataset [30].

To evaluate how our framework compares to *E. coli* experimental data, we examined LK-DFBA (NLR), as it was the best approach in the case of low sampling frequency and high noise (Figure 3B). Figure 5 shows the average Pearson correlation of the LK-DFBA (NLR) flux predictions (after being fitted to ten noisy datasets with nT = 15 and CoV = 0.15) and the average correlation of the ODE model flux predictions with the experimental steady-state flux data. Of the gene knockouts and fluxes examined, LK-DFBA (NLR) generally gave reliable predictions for whether fluxes increased or decreased due to gene knockouts, with correlation values greater than 0.6 in all but two cases and correlations greater than 0.7 in 6 out of 13 cases. These correlations were very similar to the correlations yielded by the ODE-based model. In 10 out of 13 knockouts, the correlations calculated for LK-DFBA outperformed or were within 5% of the correlations calculated with the ODE-based model. These results support the significant promise of LK-DFBA approaches for predictivity comparable to that of standard models but with the additional benefits (including relative model simplicity and potential scalability) that accrue from using a LP-based formulation.

**Figure 5:**
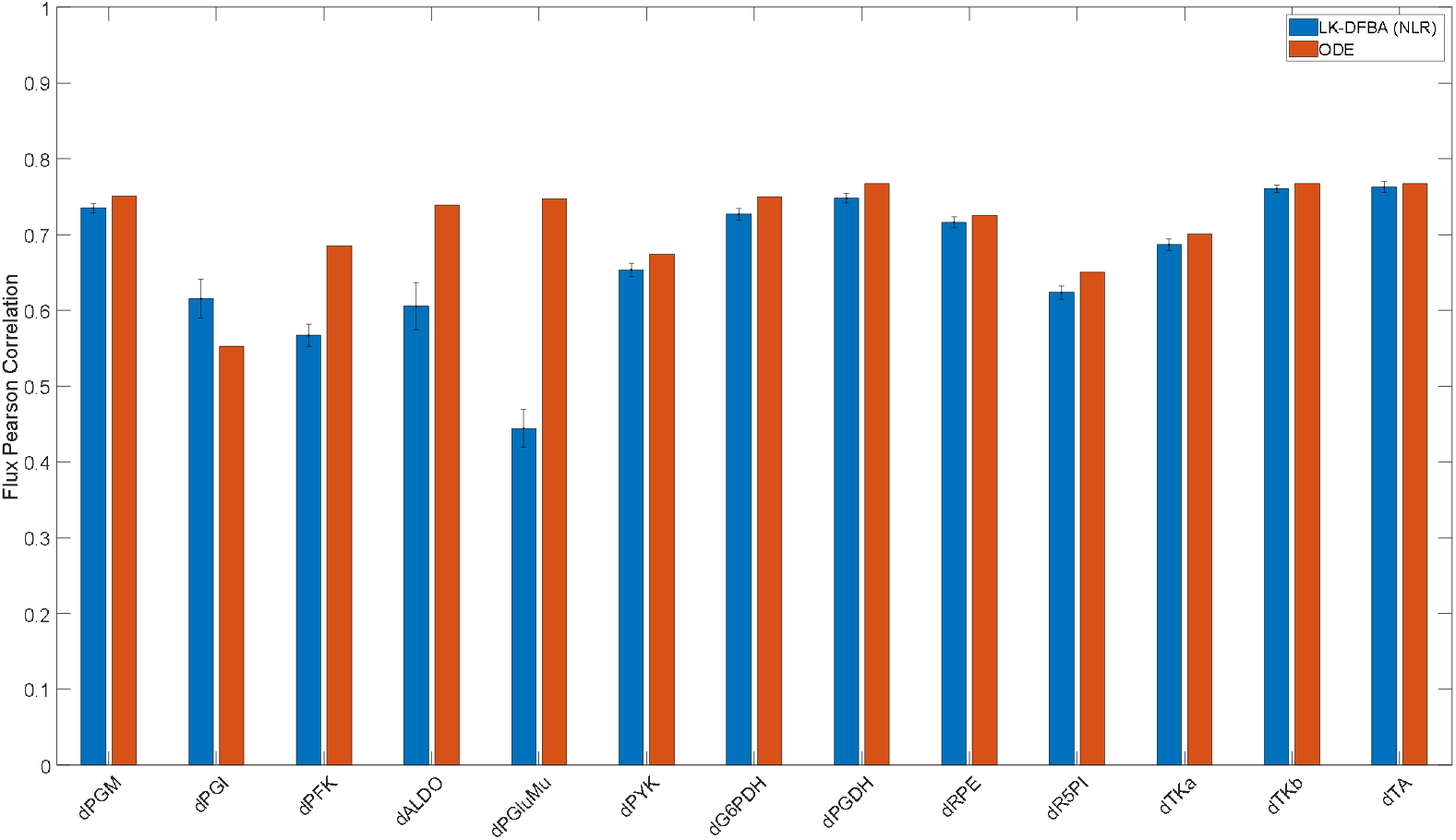
Pearson correlation coefficients of LK-DFBA and ODE model flux predictions with *E. coli* experimental data. LK-DFBA (NLR) was the best approach when fitting on low sampling frequency (nT = 15) and high noise (CoV = 0.15) data. Blue and red bars represent LK-DFBA (NLR) and ODE model mean correlations, respectively, between the average predicted flux profiles and experimental steady-state flux data for various gene knockout conditions. Gene knockouts in the LK-DFBA and ODE-based models were simulated by down-regulating relevant pathways. Error bars for LK-DFBA represent one standard deviation (N = 10 runs).

## Discussion

At the outset of performing this work, we intended to identify an LK-DFBA constraint approach to improve upon the originally published framework that used only linear kinetics constraints. Instead, we have demonstrated that the best constraint approach depends on both the topology and the parameters of the system being modeled. Despite each of the 20 small synthetic models having the exact same stoichiometric and regulatory topology, the optimal LK-DFBA approach varied across these 20 models, and for the majority of the models one of the new constraint approaches performed the best. This finding suggests that the emergent dynamics from the collective metabolic reactions are just as important as the topology of the system in determining which constraint approach best fits data from the system. It also supports the potential benefit of having multiple types of constraints to choose from, as done in this work, to enable more accurate modeling of any given system.

These findings from synthetic systems were reinforced by analysis of biological systems based on *E. coli* and *L. lactis* metabolism. While LK-DFBA (HP) performed the best on *L. lactis* noiseless data, LK-DFBA (NLR) performed the best on *E. coli* noiseless data. (We do note, though, that LK-DFBA (NLR) was superior for both systems under the lowest sampling frequency and highest noise data conditions). We further confirmed that topology is not the sole factor that determines the optimal constraint approach for a given system by randomizing parameters in the *E. coli* model (Figure S8): again, the best constraint approach varied across these topologically identical new models. This finding has particular relevance to the development of metabolic models for organisms beyond those few that are widely studied: many metabolic pathways are topologically well-conserved across many species (e.g. central carbon metabolism [1]), though the kinetic and regulatory parameters in these pathways can be vastly different. Thus, having multiple constraint approaches to choose from will improve the ability to model different systems using LK-DFBA.

Critically, the potential use of multiple types of constraints for a given model does not appear to contribute substantially to overfitting. While the best constraint approach varied across different model parameterizations and topologies, the best approach for a given model for predicting metabolic phenotypes due to genetic perturbations was generally consistent across a wide range of perturbations. This trend remained true whether using noiseless data, data with low sampling frequency and high noise, or noisy data that had been smoothed. These results instill confidence that the best constraint approach found when fitting to a wild-type metabolic system will also be the best approach when predicting changes to that system, meaning that an approach entailing the selection of the best-fitting of multiple constraint frameworks is viable, likely to be successful, and unlikely to contribute substantially to overfitting. We speculate that the reason for the success of this approach may be due to the robustness of the emergent metabolic dynamics of a system. Resampling all of the kinetic parameters in a system (as we did for 20 synthetic models and 5 *E. coli* models) can yield qualitatively different metabolite or flux profiles exhibiting qualitatively different behaviors that are better captured by different types of constraints. When individual genes or pathways are down- or up-regulated, though, it is common for only the nearest neighboring metabolites or pathways to be significantly affected, meaning that the emergent behavior from the system would not change too greatly and thus the same constraint approach would be optimal. To easily construct the optimal LK-DFBA model for a given biological system, we envision the workflow presented in Figure 6. After compiling the relevant system stoichiometry, regulatory structure, and metabolomics and fluxomics data, one can fit each of the four LK-DFBA approaches to the data and determine which constraint approach is optimal. Again, based on our findings, this optimal constraint approach fitted to the wild-type dataset is the most likely to work the best for predicting the results of model perturbations.

**Figure 6:**
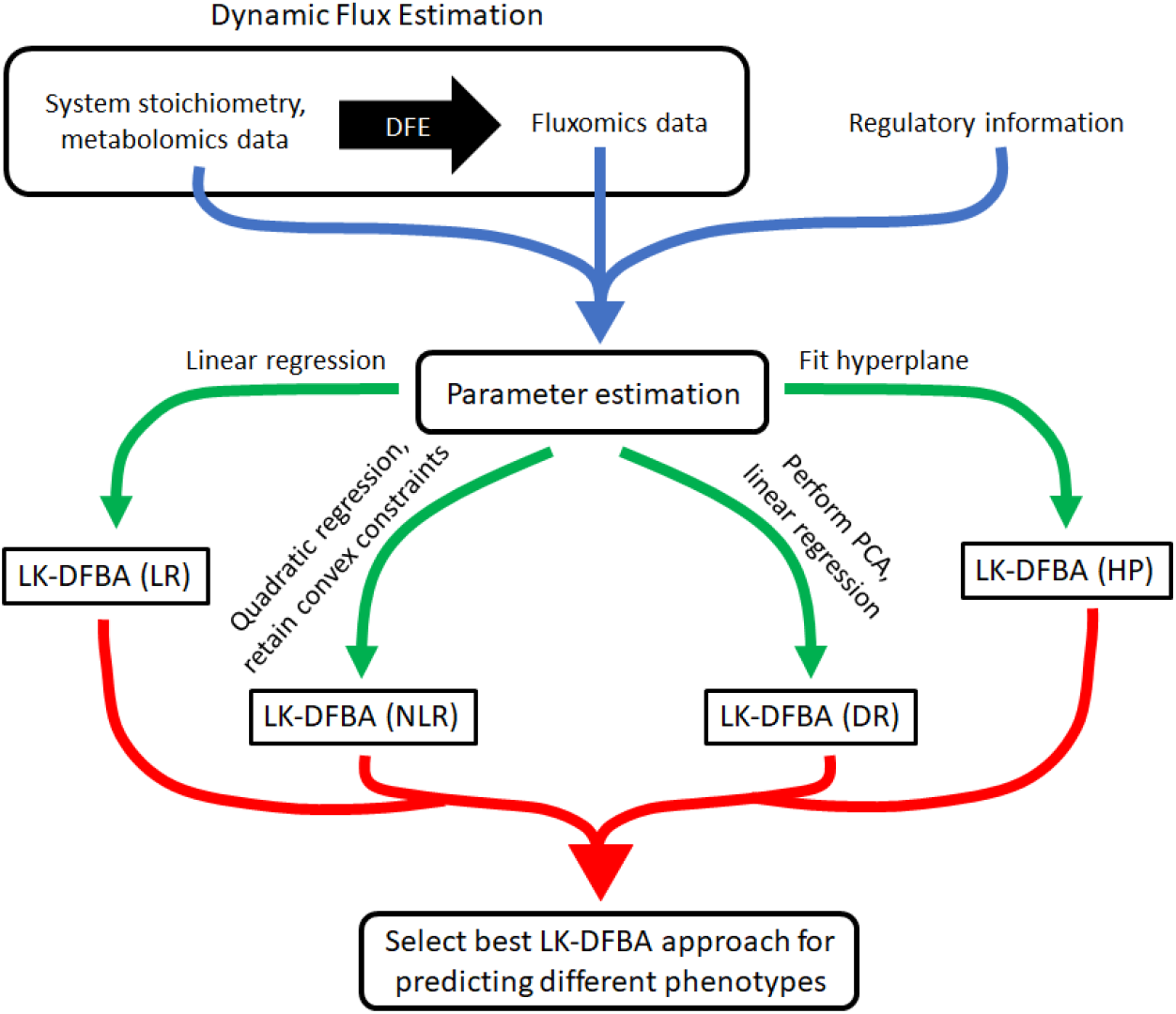
Workflow for selecting the best constraint approach for LK-DFBA when modeling metabolic systems. Dynamic Flux Estimation (DFE) is applied to the system stoichiometry and available metabolomics data to infer instantaneous fluxes. The system stoichiometry, metabolomics data, inferred flux data, and system regulatory information are then used to estimate parameters for each LK-DFBA approach (blue arrow). Using multiple constraint approaches (green arrows), four different LK-DFBA models are created and tested for their respective abilities to recapitulate training data. The model with the lowest error is selected and can be used for future *in silico* predictions (red arrow).

Beyond these investigations into the utility and robustness of new types of constraints for LK-DFBA, we also compared LK-DFBA predictions to experimental data. Using ODE models and experimental data from *L. lactis* and *E. coli*, we found that LK-DFBA can effectively predict qualitative trends in concentration profiles of some important metabolites. While we have previously shown that LK-DFBA captures metabolite dynamics in synthetic data generated by ODE models, this is the first time LK-DFBA predictions have been validated with experimental data. For key metabolites that are important inputs and outputs of the system (e.g. carbon sources or end products), LK-DFBA can qualitatively predict if their concentration profiles are expected to decrease or increase, which is an important capability if one is using LK-DFBA to engineer organisms to efficiently produce certain metabolites. Cofactors, on the other hand, are more difficult to model using LK-DFBA but are still typically predicted to be within an order of magnitude of the experimental data in most cases. This capability could be useful when assessing levels of accumulating toxic metabolites or cofactor imbalances if exact concentrations are not necessary. We also found that LK-DFBA flux profile predictions were highly correlated with experimental flux data from genetic knockout experiments. Furthermore, these correlations were comparable to those found when using the ODE-based model. We note, though, that this comparable predictivity is limited by the fact that LK-DFBA was trained using ODE-generated data; if it had instead been fitted to actual metabolomics and fluxomics time course data used in the Ishii experiments (which is not available), these correlation values could possibly be even higher. Similarly, an improved objective function over the reasonable but arbitrary and unoptimized one used here could also lead to significant improvements in the performance of LK-DFBA.

By showing for the first time that LK-DFBA can predict changes in metabolite concentrations and flux profiles qualitatively, these results support LK-DFBA’s potential as a widely-applicable metabolic modeling tool. While many ODE-based modeling approaches require specific kinetic equations for each flux reaction, LK-DFBA is more generalized. With four types of kinetics constraints that account for different biological interaction phenomena between metabolites and fluxes, we have made LK-DFBA more amenable to different systems. Additionally, applying the four LK-DFBA approaches to these models of *L. lactis* and *E. coli* has established that our framework can handle various biological systems of substantial size without the need for computationally taxing parameter estimation steps (using only regression for parameter estimation, as in this work). Because each of the four LK-DFBA approaches maintains an easily solvable LP or QCP structure, LK-DFBA is a prime candidate for scaling up to model a variety of genome-scale systems while also capturing their metabolite dynamics.

While the addition of new constraint approaches has significantly improved the original LK-DFBA (LR) framework, there are still several areas where LK-DFBA can be improved. If computational resources when building the model are not a concern, a secondary optimization step can be used, as in the LK-DFBA (LR+) approach, to improve the parameters in each of the new constraint approaches. In addition, as previously noted the objective function used in LK-DFBA is also a ripe target for future efforts to improve this modeling framework. Here we have chosen objective functions that lead to the maximization of putatively important fluxes, but unlike many other constraint-based models, there was no specific biomass or other objective flux to use. Optimizing the weight of each flux or metabolite in the objective function could lead to even lower observed errors compared to experimental data and may also provide insight into the selective pressures on real biological systems.

## Conclusion

In this work, we have shown that the LK-DFBA modeling framework can be improved by implementing more complex constraints with increased biological relevance. We showed that there is no single best LK-DFBA constraint approach for all models, and that the optimal approach depends not just on the topology of the biochemical system but also its kinetics and parameters. Importantly, though, the constraint approach that performs the best in recapitulating training data consistently outperforms other constraint approaches at predicting the results of metabolic perturbations on the same system, suggesting that selecting between multiple different types of constraints improves model predictivity with limited contributions to overfitting. With these new constraint approaches, we are able to model a variety of metabolic systems more accurately than if we were to just use the original LK-DFBA (LR) method. Moreover, based on comparisons to experimental data, we showed that the improved LK-DFBA approaches can reasonably capture the qualitative dynamics of important metabolites and fluxes of interest to researchers. While these predictions may not be smooth or quantitative, the qualitative prediction of trends of metabolite dynamics in response to major perturbations is arguably the most critical aspect needed for applications like metabolic engineering: knowing that a specific knockout will increase or decrease flux is often sufficient to justify the expense of experimental implementation of such genetic perturbations. Moreover, we expect this computational framework to (with future effort) provide the potential for computationally reasonable scale-up to the genome scale. While the acquisition of quality metabolomics and fluxomics data to build the constraints in LK-DFBA is still a challenge, the work we have presented here lays the groundwork needed to take full advantage of these types of datasets as they become increasingly more readily available.

## Supporting information

Ecoli parameters

## Data availability

The code and datasets generated during the current study are available at https://github.com/gtstylab/lk-dfba-constraints.

## Author contributions

JYL conceived of the study, participated in the design of the study, carried out computational experiments, analyzed experimental results, and helped to draft the manuscript. MPS conceived of the study, participated in the design of the study, analyzed experimental results, and helped to draft the manuscript. All authors read and approved of the final manuscript.

