## Supplementary for "Diverse classes of constraints enable broader applicability of a linear programming-based dynamic metabolic modeling framework"

### Methods

**Table S1: Kinetic parameters in synthetic models** Parameters were randomly generated using a uniform distribution between 0.1 and 1.0 (or -1.0 to -0.1 for $b_{2r4}$, which is a parameter that describes inhibition in the system). *k* represents the model number.

| | Kinetics | | | | | | | | | | | Initial Conditions | | | | |
|---|---|---|---|---|---|---|---|---|---|---|---|---|---|---|---|---|
| *k* | $a_2$ | $b_{21}$ | $b_{2r4}$ | $a_3$ | $b_{32}$ | $a_4$ | $b_{42}$ | $b_{4r3}$ | $a_5$ | $b_{53}$ | $b_{54}$ | $X_1$ | $X_2$ | $X_3$ | $X_4$ | $X_{BM}$ |
| 1 | 0.8 | 0.5 | -0.2 | 1.0 | 0.75 | 0.5 | 0.4 | 0.8 | 0.5 | 0.5 | 0.8 | 0.1 | 0.2 | 0.3 | 0.4 | 0.5 |
| 2 | 0.7 | 0.5 | -0.2 | 0.8 | 0.2 | 0.2 | 0.9 | 0.7 | 0.9 | 0.4 | 0.9 | 0.3 | 0.3 | 0.6 | 0.5 | 0.2 |
| 3 | 0.1 | 0.5 | -0.7 | 0.6 | 0.9 | 0.6 | 0.3 | 0.9 | 0.2 | 0.4 | 0.3 | 0.4 | 0.9 | 0.6 | 0.1 | 0.9 |
| 4 | 0.9 | 0.4 | -0.2 | 0.6 | 0.8 | 0.4 | 0.9 | 0.6 | 0.5 | 0.2 | 0.7 | 0.8 | 0.1 | 0.5 | 0.7 | 0.8 |
| 5 | 0.2 | 0.4 | -0.2 | 0.9 | 0.6 | 0.6 | 0.3 | 0.6 | 0.5 | 0.7 | 0.8 | 0.5 | 0.5 | 0.8 | 0.5 | 0.2 |
| 6 | 0.1 | 0.6 | -0.3 | 0.3 | 0.9 | 0.5 | 0.1 | 0.7 | 0.6 | 0.8 | 0.6 | 0.2 | 0.9 | 0.2 | 0.1 | 0.3 |
| 7 | 0.5 | 0.2 | -0.2 | 0.1 | 0.9 | 0.2 | 0.9 | 0.9 | 0.6 | 0.5 | 0.8 | 0.2 | 0.3 | 0.1 | 0.7 | 0.6 |
| 8 | 0.9 | 0.4 | -0.3 | 0.8 | 0.1 | 0.2 | 0.9 | 0.7 | 0.2 | 0.6 | 0.8 | 0.6 | 0.4 | 0.1 | 0.4 | 0.4 |
| 9 | 0.6 | 0.5 | -0.9 | 0.4 | 0.6 | 0.8 | 0.8 | 0.4 | 0.7 | 0.8 | 0.9 | 0.3 | 0.4 | 0.2 | 0.7 | 0.8 |
| 10 | 0.6 | 0.3 | -0.9 | 0.3 | 0.2 | 0.6 | 0.6 | 0.7 | 0.6 | 0.8 | 0.9 | 0.4 | 0.2 | 0.8 | 0.1 | 0.7 |
| 11 | 0.8 | 0.2 | -0.6 | 0.7 | 0.4 | 0.3 | 0.7 | 0.8 | 0.4 | 0.1 | 0.2 | 0.5 | 0.1 | 0.4 | 0.8 | 0.4 |
| 12 | 0.9 | 0.9 | -0.1 | 0.1 | 0.4 | 0.7 | 0.3 | 0.3 | 0.8 | 0.1 | 0.8 | 0.1 | 0.9 | 0.9 | 0.6 | 0.1 |
| 13 | 0.2 | 0.9 | -0.8 | 0.3 | 0.7 | 0.6 | 0.5 | 0.8 | 0.6 | 0.5 | 0.3 | 0.3 | 0.1 | 0.2 | 0.4 | 0.9 |
| 14 | 0.9 | 0.5 | -0.9 | 0.1 | 0.8 | 0.2 | 0.7 | 0.3 | 0.5 | 0.8 | 0.5 | 0.9 | 0.7 | 0.3 | 0.5 | 0.9 |
| 15 | 0.6 | 0.8 | -0.7 | 0.1 | 0.2 | 0.2 | 0.9 | 0.9 | 0.9 | 0.9 | 0.2 | 0.2 | 0.8 | 0.2 | 0.4 | 0.5 |
| 16 | 0.1 | 0.2 | -0.7 | 0.8 | 0.5 | 0.5 | 0.9 | 0.4 | 0.3 | 0.2 | 0.6 | 0.8 | 0.8 | 0.2 | 0.1 | 0.5 |
| 17 | 0.3 | 0.4 | -0.3 | 0.7 | 0.5 | 0.9 | 0.5 | 0.2 | 0.7 | 0.6 | 0.3 | 0.5 | 0.1 | 0.8 | 0.3 | 0.2 |
| 18 | 0.5 | 0.9 | -0.4 | 0.3 | 0.6 | 0.4 | 0.2 | 0.3 | 0.7 | 0.5 | 0.6 | 0.9 | 0.4 | 0.6 | 0.2 | 0.9 |
| 19 | 0.9 | 0.8 | -0.6 | 0.9 | 0.7 | 0.6 | 0.2 | 0.6 | 0.4 | 0.1 | 0.7 | 0.1 | 0.3 | 0.5 | 0.2 | 0.2 |
| 20 | 0.9 | 0.9 | -0.2 | 0.1 | 0.7 | 0.3 | 0.3 | 0.5 | 0.6 | 0.4 | 0.7 | 0.4 | 0.8 | 0.2 | 0.3 | 0.4 |

#### Kinetic parameters in *E. coli* models

Parameters were randomly generated by drawing from the random normal distribution $N_i \sim (p_i, p_i)$, where $p_i$ is the original value of the *i*th parameter. If the random parameter was negative, the absolute value of the parameter was used. Some parameters, such as the feed rate, dilution rate, and rates of synthesis reactions were kept at their original parameter values to ensure the models were viable. The parameters for each model can be found in ecoli_parameters.xlsx.

#### LK-DFBA Objective Functions

Like other constraint-based methods, LK-DFBA requires an objective function, which is usually tied to some presumed goal of the system (such as maximizing biomass or ATP production) that stems from evolutionary pressure. FBA models for specific organisms commonly have a separate flux reaction dedicated to biomass, made up of precise ratios of different metabolites. While LK-DFBA models with tuned objective functions can be created, the biological models we sought to use here do not have pre-existing tuned objective functions, so we instead focused on LK-DFBA's performance using generic objective functions.

Here, we have chosen flux $v_5$ as the objective function for the synthetic model, as it is the only efflux out of the system. For the *L. lactis* model, we use the LDH pathway as the objective function to maximize production of lactate because it is a key metabolite in the organism (which is commonly used for dairy products) and was the metabolite produced at the highest levels in the original *L. lactis* model [1]. The objective function used for the *E. coli* model was to maximize all effluxes from the system, which included murein synthesis, glycerol-3-phosphate dehydrogenase, serine synthesis, PEP carboxylase, DAHP synthesis, pyruvate dehydrogenase, ribose phosphate pyrophosphokinase, glucose-1-phosphate adenyltransferase, the synthesis of murein and chorismate from PEP, and the synthesis of isoleucine, alanine, α-ketoisovalerate, and diaminopimelate from pyruvate. While we have observed that these objective functions can be further improved, and approaches have been developed for finding an optimal objective function for a model by creating a bilevel optimization problem and then leveraging the duality theorem [2, 3], our chosen objective functions were sufficient to at least qualitatively model the synthetic, *L. lactis*, and *E. coli* systems.

**Translating constraints to contain training data**

We found that translating the constraints such that all training data fall in the region under the inequality constraint decreased the possibility of the computational solver encountering infeasible solutions when simulating metabolite dynamics. Thus, for all LK-DFBA approaches, each constraint was translated to contain the training data by increasing the intercept of the constraint (i.e. the *b* parameter in LK-DFBA (LR), LK-DFBA (DR), and LK-DFBA (HP), and the *c* parameter in LK-DFBA (NLR)) until no training data were above each constraint.

**Error Calculation**

The error of the predictions made by LK-DFBA was calculated using a normalized root mean squared error (NRMSE) between the LK-DFBA predicted metabolite concentrations and the noiseless ODE concentration or experimental data. $P_{ik}$ and $R_{ik}$ are the predicted (e.g. results from LK-DFBA) and reference (e.g. ODE model or experimental) data from a system with $m$ metabolites and $nT$ time points. $\bar{R}_i$ is the mean of the concentrations of reference metabolite $i$ across all time points to normalize the data and $N$ is the total number of data points used in the NRMSE calculation.

$$\boldsymbol{NRMSE} = \sqrt{\frac{\sum_{i=1}^{m}\sum_{k=1}^{nT}\left(\frac{P_{ik}-R_{ik}}{\bar{R}_i}\right)^2}{N}} \quad (2)$$

**Pearson Correlation Calculation**

The available *E. coli* knockout experimental data consisted of steady-state flux data, so to compare these to the knockout predictions made by LK-DFBA (which did not yield a steady state over the ten second time interval of the model) we used the average flux of our time course predictions. Because the average flux of our predictions and the steady-state fluxes of the experimental data are different measurements and therefore not directly comparable using NRMSE, we chose to use a Pearson correlation coefficient to evaluate our framework, which was recently used in a similar comparative analysis of metabolic models [4]. High correlations between steady-state flux experimental data and the average flux predictions would indicate that LK-DFBA can effectively predict if gene knockouts lead to an increase or decrease in flux for modeled reactions. The calculation for the Pearson correlation coefficient is shown below, where $A_i$ is the average of the predicted flux profile for the *i*th flux, $v_i$ is the flux value of the *i*th flux from the experimental data, $\bar{A}$ is the mean across all fluxes for the average of computationally predicted fluxes, $\bar{v}$ is the mean flux value across all fluxes for the experimental data, and $n$ is double the number of fluxes that are shared between both the *E. coli* model and experimental data because it includes flux values before and after the gene knockout ($n$ = 28).

$$\boldsymbol{Pearson\ Correlation\ Coefficient} = \frac{\sum_{i=1}^{n}(A_i-\bar{A})(v_i-\bar{v})}{\sqrt{\sum_{i=1}^{n}(A_i-\bar{A})^2\sum_{i=1}^{n}(v_i-\bar{v})^2}} \tag{3}$$

| Method | Phenotype | Model # 1 | 2 | 3 | 4 | 5 | 6 | 7 | 8 | 9 | 10 | 11 | 12 | 13 | 14 | 15 | 16 | 17 | 18 | 19 | 20 |
|---|---|---|---|---|---|---|---|---|---|---|---|---|---|---|---|---|---|---|---|---|---|
| LK-DFBA (LR) | WT | 0.10 | 0.17 | 0.93 | 0.05 | 1.07 | 0.43 | 0.10 | 0.46 | 0.18 | 1.01 | 0.49 | 0.17 | 0.17 | 0.54 | 0.21 | 0.53 | 0.76 | 0.13 | 0.08 | 0.05 |
| | dV2 | 0.44 | 0.34 | 0.57 | 0.45 | 1.85 | 0.52 | 0.46 | 0.67 | 0.61 | 2.27 | 0.67 | 0.99 | 0.44 | 0.52 | 0.38 | 0.57 | 0.64 | 0.47 | 0.51 | 0.45 |
| | uV2 | 0.47 | 0.28 | 1.56 | 0.41 | 0.75 | 0.30 | 0.56 | 0.44 | 0.83 | 0.64 | 0.56 | 0.59 | 0.72 | 0.52 | 0.48 | 1.44 | 0.75 | 0.51 | 0.48 | 0.40 |
| | dV3 | 0.17 | 0.49 | 1.14 | 0.34 | 0.68 | 0.63 | 0.29 | 0.63 | 0.51 | 1.01 | 0.60 | 1.91 | 0.30 | 0.65 | 0.52 | 0.92 | 0.49 | 0.31 | 0.11 | 0.69 |
| | uV3 | 0.47 | 2.29 | 0.92 | 0.39 | 1.80 | 0.32 | 0.25 | 1.59 | 0.37 | 0.98 | 0.57 | 0.86 | 0.46 | 0.64 | 0.40 | 0.79 | 1.60 | 0.63 | 0.21 | 0.36 |
| | dV4 | 0.54 | 0.30 | 0.63 | 0.28 | 0.57 | 0.50 | 0.27 | 0.51 | 0.17 | 0.74 | 0.56 | 0.38 | 0.25 | 0.49 | 0.12 | 0.87 | 0.67 | 0.22 | 0.64 | 0.20 |
| | uV4 | 0.35 | 0.22 | 1.72 | 0.35 | 1.59 | 0.77 | 0.30 | 0.63 | 0.58 | 2.57 | 0.64 | 1.49 | 0.56 | 0.85 | 0.42 | 0.73 | 0.87 | 0.67 | 0.49 | 0.86 |
| LK-DFBA (NLR) | WT | 0.09 | 0.11 | 0.89 | 0.07 | 1.03 | 0.33 | 0.13 | 0.36 | 0.14 | 0.47 | 0.33 | 0.14 | 0.22 | 0.45 | 0.20 | 0.67 | 0.34 | 0.12 | 0.11 | 0.02 |
| | dV2 | 0.33 | 0.48 | 0.44 | 0.31 | 2.07 | 0.33 | 0.45 | 0.57 | 0.26 | 1.12 | 0.53 | 1.83 | 0.38 | 0.32 | 0.35 | 0.26 | 0.86 | 0.31 | 0.36 | 0.35 |
| | uV2 | 0.23 | 0.14 | 1.70 | 0.34 | 1.29 | 0.51 | 0.20 | 0.59 | 0.18 | 0.19 | 0.30 | 0.40 | 0.46 | 0.35 | 0.33 | 1.14 | 0.28 | 0.15 | 0.24 | 0.10 |
| | dV3 | 0.23 | 0.39 | 1.17 | 0.37 | 0.58 | 0.41 | 0.35 | 0.51 | 0.33 | 0.44 | 0.33 | 3.28 | 0.43 | 0.65 | 1.05 | 0.93 | 0.59 | 0.49 | 0.19 | 0.54 |
| | uV3 | 0.49 | 2.25 | 0.99 | 0.40 | 2.02 | 0.32 | 0.24 | 1.72 | 0.39 | 1.78 | 0.47 | 0.83 | 0.53 | 0.55 | 0.38 | 1.22 | 1.02 | 0.65 | 0.15 | 0.36 |
| | dV4 | 0.44 | 0.30 | 0.44 | 0.47 | 0.52 | 0.26 | 0.24 | 0.43 | 0.31 | 0.19 | 0.40 | 0.37 | 0.31 | 0.23 | 0.09 | 0.61 | 0.42 | 0.40 | 0.57 | 0.24 |
| | uV4 | 0.34 | 0.22 | 1.74 | 0.35 | 1.41 | 0.73 | 0.27 | 0.61 | 0.54 | 1.29 | 0.57 | 1.26 | 0.59 | 0.72 | 0.40 | 1.25 | 0.92 | 0.81 | 0.51 | 0.66 |
| LK-DFBA (DR) | WT | 0.73 | 0.39 | 0.33 | 0.47 | 0.35 | 0.20 | 0.29 | 0.83 | 0.14 | 1.67 | 0.94 | 0.17 | 0.42 | 0.66 | 0.24 | 0.72 | 0.34 | 0.23 | 0.28 | 0.17 |
| | dV2 | 0.40 | 0.46 | 0.21 | 0.41 | 0.43 | 0.39 | 0.46 | 0.73 | 0.47 | 2.94 | 0.61 | 0.28 | 0.34 | 0.63 | 0.36 | 0.57 | 0.74 | 0.36 | 0.49 | 0.34 |
| | uV2 | 0.82 | 0.50 | 0.67 | 0.70 | 0.13 | 0.21 | 0.49 | 0.80 | 0.57 | 1.28 | 0.83 | 0.29 | 0.76 | 0.64 | 0.48 | 1.03 | 0.64 | 0.48 | 0.52 | 0.45 |
| | dV3 | 0.70 | 0.61 | 0.39 | 0.65 | 0.21 | 0.37 | 0.39 | 0.85 | 0.51 | 1.97 | 0.92 | 0.20 | 0.59 | 0.70 | 0.53 | 1.43 | 0.37 | 0.50 | 0.29 | 1.11 |
| | uV3 | 0.91 | 0.75 | 0.33 | 0.36 | 0.56 | 0.17 | 0.50 | 0.82 | 0.71 | 1.18 | 0.69 | 1.65 | 0.79 | 0.71 | 0.62 | 0.42 | 1.07 | 0.28 | 0.36 | 0.33 |
| | dV4 | 0.46 | 0.76 | 0.20 | 0.26 | 0.21 | 0.11 | 0.22 | 0.51 | 0.46 | 0.92 | 0.68 | 0.79 | 0.17 | 0.52 | 0.14 | 0.29 | 0.43 | 0.14 | 0.18 | 0.11 |
| | uV4 | 0.70 | 0.52 | 0.45 | 1.03 | 0.48 | 0.49 | 0.74 | 1.29 | 0.53 | 4.33 | 1.56 | 0.22 | 0.68 | 1.15 | 0.42 | 1.39 | 0.82 | 2.23 | 0.47 | 1.87 |
| LK-DFBA (HP) | WT | 0.01 | 1.32 | 1.95 | 0.01 | 1.63 | 1.01 | 0.43 | 0.03 | 2.67 | 2.31 | 1.06 | 2.17 | 1.49 | 0.89 | 0.54 | 1.11 | 0.44 | 1.41 | 1.24 | 1.14 |
| | dV2 | 0.34 | 2.31 | 1.49 | 0.57 | 0.94 | 1.15 | 0.23 | 2.80 | 1.31 | 4.10 | 0.73 | 3.69 | 1.17 | 0.93 | 0.44 | 1.50 | 0.83 | 1.43 | 1.53 | 1.19 |
| | uV2 | 0.43 | 1.35 | 2.48 | 0.18 | 2.05 | 1.54 | 0.74 | 0.27 | 3.95 | 1.73 | 0.86 | 2.55 | 1.67 | 0.82 | 0.61 | 0.58 | 0.54 | 1.70 | 1.07 | 1.16 |
| | dV3 | 0.20 | 0.52 | 2.30 | 0.03 | 0.79 | 0.89 | 0.76 | 0.21 | 2.94 | 1.80 | 1.09 | 6.54 | 1.23 | 0.86 | 0.29 | 0.75 | 0.55 | 1.64 | 1.36 | 1.33 |
| | uV3 | 0.11 | 8.71 | 1.53 | 0.88 | 3.83 | 1.20 | 0.52 | 1.06 | 1.92 | 2.30 | 0.78 | 1.20 | 1.44 | 0.86 | 0.68 | 1.98 | 0.95 | 1.32 | 1.20 | 0.59 |
| | dV4 | 0.05 | 1.59 | 1.04 | 0.13 | 1.93 | 0.76 | 0.23 | 0.08 | 1.38 | 1.08 | 0.70 | 1.3 | 0.82 | 0.64 | 0.43 | 1.60 | 0.73 | 0.82 | 0.73 | 0.64 |
| | uV4 | 0.10 | 1.07 | 3.45 | 0.06 | 1.40 | 1.70 | 0.82 | 0.56 | 5.17 | 4.83 | 1.77 | 7.70 | 2.69 | 1.48 | 0.66 | 0.86 | 0.87 | 3.60 | 2.67 | 1.94 |

**Figure S1: LK-DFBA performance on noiseless synthetic model data** Each constraint approach was used to fit parameters to noiseless (nT = 50) wild-type (WT) data and then used to simulate the WT system and the system with *in silico* genetic perturbations with fluxes $v_2$, $v_3$, or $v_4$ down- or up-regulated. Dark green boxes represent the lowest NRMSE within each phenotype for each synthetic model, while dark red boxes represent the highest NRMSE. The cells with bolded white numbers indicate the LK-DFBA approach that best fits the WT data (also highlighted in cyan). Cells with white numbers are generally consistently green, indicating that fitting to WT data is a good indicator of which approach will be optimal across perturbations.

| Approach | Phenotype | Model # 1 | 2 | 3 | 4 | 5 | 6 | 7 | 8 | 9 | 10 | 11 | 12 | 13 | 14 | 15 | 16 | 17 | 18 | 19 | 20 |
|---|---|---|---|---|---|---|---|---|---|---|---|---|---|---|---|---|---|---|---|---|---|
| LK-DFBA (LR) | WT | 0.11 | 0.17 | 0.94 | 0.09 | 1.06 | 0.38 | 0.10 | 0.35 | 0.20 | 1.08 | 0.49 | 0.19 | 0.17 | 0.54 | 0.20 | 0.52 | 0.82 | 0.20 | 0.10 | 0.05 |
| | dV2 | 0.43 | 0.36 | 0.74 | 0.47 | 1.54 | 0.50 | 0.42 | 0.62 | 0.65 | 2.11 | 0.67 | 0.91 | 0.45 | 0.52 | 0.37 | 0.57 | 0.69 | 0.48 | 0.56 | 0.45 |
| | uV2 | 0.47 | 0.27 | 1.33 | 0.40 | 0.84 | 0.37 | 0.54 | 0.43 | 0.86 | 0.63 | 0.56 | 0.46 | 0.72 | 0.52 | 0.48 | 0.61 | 0.79 | 0.45 | 0.41 | 0.37 |
| | dV3 | 0.18 | 0.48 | 1.04 | 0.37 | 0.59 | 0.55 | 0.31 | 0.63 | 0.64 | 0.99 | 0.61 | 1.98 | 0.31 | 0.65 | 0.50 | 0.84 | 0.51 | 0.24 | 0.19 | 0.67 |
| | uV3 | 0.47 | 2.26 | 1.06 | 0.41 | 1.85 | 0.32 | 0.25 | 1.58 | 0.36 | 1.06 | 0.59 | 0.87 | 0.46 | 0.63 | 0.39 | 0.77 | 1.70 | 0.65 | 0.38 | 0.36 |
| | dV4 | 0.51 | 0.30 | 0.69 | 0.29 | 0.54 | 0.46 | 0.26 | 0.50 | 0.18 | 0.64 | 0.59 | 0.39 | 0.24 | 0.49 | 0.11 | 0.83 | 0.75 | 0.22 | 0.70 | 0.20 |
| | uV4 | 0.35 | 0.22 | 1.55 | 0.37 | 1.12 | 0.72 | 0.29 | 0.59 | 0.60 | 2.51 | 0.65 | 1.60 | 0.56 | 0.82 | 0.40 | 0.69 | 0.89 | 0.62 | 0.49 | 0.79 |
| LK-DFBA (NLR) | WT | 0.08 | 0.13 | 1.23 | 0.12 | 1.03 | 0.31 | 0.15 | 0.22 | 0.16 | 0.32 | 0.32 | 0.20 | 0.21 | 0.41 | 0.17 | 0.43 | 0.36 | 0.14 | 0.19 | 0.08 |
| | dV2 | 0.33 | 0.45 | 0.63 | 0.37 | 2.39 | 0.32 | 0.42 | 0.57 | 0.26 | 0.83 | 0.53 | 1.71 | 0.38 | 0.31 | 0.32 | 0.23 | 0.71 | 0.32 | 0.41 | 0.38 |
| | uV2 | 0.11 | 0.19 | 1.50 | 0.29 | 1.12 | 0.50 | 0.18 | 0.61 | 0.17 | 0.22 | 0.31 | 0.42 | 0.34 | 0.21 | 0.28 | 0.90 | 0.30 | 0.18 | 0.31 | 0.10 |
| | dV3 | 0.34 | 0.42 | 1.08 | 0.32 | 0.49 | 0.35 | 0.34 | 0.61 | 0.35 | 0.40 | 0.36 | 3.06 | 0.34 | 0.58 | 0.85 | 0.82 | 0.46 | 0.50 | 0.22 | 0.56 |
| | uV3 | 0.48 | 2.26 | 1.74 | 0.37 | 2.48 | 0.32 | 0.25 | 1.68 | 0.40 | 0.87 | 0.49 | 0.84 | 0.51 | 0.48 | 0.37 | 0.81 | 0.97 | 0.62 | 0.44 | 0.37 |
| | dV4 | 0.19 | 0.30 | 0.68 | 0.34 | 0.55 | 0.25 | 0.25 | 0.54 | 0.30 | 0.22 | 0.40 | 0.38 | 0.26 | 0.21 | 0.10 | 0.51 | 0.42 | 0.50 | 0.57 | 0.23 |
| | uV4 | 0.42 | 0.24 | 2.27 | 0.35 | 1.55 | 0.71 | 0.29 | 0.50 | 0.60 | 0.93 | 0.51 | 0.85 | 0.57 | 0.70 | 0.38 | 0.85 | 1.01 | 0.74 | 0.56 | 0.59 |
| LK-DFBA (DR) | WT | 0.73 | 0.39 | 0.39 | 0.34 | 0.32 | 0.20 | 0.35 | 0.80 | 0.12 | 1.61 | 0.93 | 0.17 | 0.41 | 0.66 | 0.24 | 0.73 | 0.30 | 0.20 | 0.34 | 0.18 |
| | dV2 | 0.41 | 0.47 | 0.31 | 0.43 | 0.40 | 0.39 | 0.48 | 0.70 | 0.48 | 2.79 | 0.61 | 0.27 | 0.34 | 0.63 | 0.35 | 0.58 | 0.75 | 0.38 | 0.51 | 0.35 |
| | uV2 | 0.82 | 0.50 | 0.68 | 0.70 | 0.13 | 0.22 | 0.50 | 0.80 | 0.58 | 1.26 | 0.83 | 0.40 | 0.76 | 0.64 | 0.48 | 1.03 | 0.76 | 0.48 | 0.52 | 0.45 |
| | dV3 | 0.71 | 0.61 | 0.44 | 0.61 | 0.27 | 0.36 | 0.33 | 0.75 | 1.38 | 1.94 | 0.91 | 0.22 | 0.57 | 0.69 | 0.53 | 1.40 | 0.39 | 0.50 | 0.36 | 1.09 |
| | uV3 | 0.92 | 0.75 | 0.40 | 0.33 | 0.49 | 0.19 | 0.46 | 0.81 | 0.72 | 1.16 | 0.69 | 1.70 | 0.78 | 0.71 | 0.62 | 0.45 | 1.09 | 0.25 | 0.43 | 0.33 |
| | dV4 | 0.46 | 0.76 | 0.29 | 0.28 | 0.18 | 0.11 | 0.19 | 0.44 | 0.43 | 0.90 | 0.68 | 0.73 | 0.17 | 0.52 | 0.13 | 0.31 | 0.41 | 0.15 | 0.17 | 0.11 |
| | uV4 | 0.72 | 0.53 | 0.54 | 0.94 | 0.49 | 0.49 | 0.69 | 1.28 | 0.53 | 4.14 | 1.55 | 0.2 | 0.53 | 1.12 | 0.40 | 1.37 | 0.82 | 2.19 | 0.55 | 1.81 |
| LK-DFBA (HP) | WT | 0.41 | 1.40 | 1.87 | 0.53 | 1.20 | 0.81 | 0.40 | 1.40 | 2.11 | 2.21 | 0.78 | 2.05 | 1.33 | 0.89 | 0.18 | 1.30 | 0.79 | 0.82 | 0.70 | 0.91 |
| | dV2 | 0.60 | 2.54 | 1.44 | 0.44 | 0.83 | 1.17 | 0.22 | 2.40 | 1.28 | 3.89 | 0.49 | 2.08 | 0.97 | 0.88 | 0.15 | 1.56 | 1.38 | 0.75 | 0.57 | 0.95 |
| | uV2 | 0.47 | 1.39 | 2.39 | 0.69 | 1.69 | 1.15 | 0.73 | 1.27 | 3.11 | 1.69 | 0.65 | 0.86 | 1.60 | 0.81 | 0.27 | 1.57 | 0.84 | 0.92 | 0.71 | 0.92 |
| | dV3 | 0.50 | 0.52 | 2.20 | 0.29 | 0.45 | 0.64 | 0.58 | 0.55 | 2.32 | 1.73 | 0.68 | 4.75 | 1.18 | 0.77 | 0.33 | 0.82 | 0.86 | 0.90 | 0.77 | 1.03 |
| | uV3 | 0.62 | 9.36 | 1.49 | 1.05 | 2.73 | 0.88 | 0.49 | 20.4 | 0.97 | 2.12 | 0.82 | 1.38 | 1.24 | 0.84 | 0.49 | 2.25 | 1.50 | 0.81 | 0.74 | 0.61 |
| | dV4 | 0.35 | 1.67 | 0.99 | 0.42 | 1.37 | 0.45 | 0.22 | 1.02 | 1.11 | 1.12 | 0.50 | 1.2 | 0.72 | 0.62 | 0.15 | 1.41 | 0.84 | 0.53 | 0.34 | 0.48 |
| | uV4 | 0.60 | 1.14 | 3.31 | 0.47 | 0.76 | 1.51 | 0.80 | 1.35 | 3.64 | 4.64 | 1.27 | 3.7 | 2.42 | 1.51 | 0.32 | 1.71 | 1.84 | 2.61 | 1.39 | 1.77 |

**Figure S2: LK-DFBA performance on noisy synthetic model data, nT = 50, CoV = 0.05** Each constraint approach was used to fit parameters to noisy (nT = 50, CoV = 0.05) wild-type (WT) data and then used to simulate the WT system and the system with *in silico* genetic perturbations with fluxes $v_2$, $v_3$, or $v_4$ down- or up-regulated. Dark green boxes represent the lowest average NRMSE (N = 10) within each phenotype for each synthetic model, while dark red boxes represent the highest average NRMSE. The cells with bolded white numbers indicate the LK-DFBA approach that best fits the WT data (also highlighted in cyan). Cells with white numbers are generally consistently green, indicating that fitting to WT data is a good indicator of which approach will be optimal across perturbations.

| | | Model # | | | | | | | | | | | | | | | | | | | |
|---|---|---|---|---|---|---|---|---|---|---|---|---|---|---|---|---|---|---|---|---|---|
| | | 1 | 2 | 3 | 4 | 5 | 6 | 7 | 8 | 9 | 10 | 11 | 12 | 13 | 14 | 15 | 16 | 17 | 18 | 19 | 20 |
| LK-DFBA (LR) | WT | 0.12 | 0.17 | 0.90 | 0.16 | 0.87 | 0.39 | 0.10 | 0.31 | 0.28 | 1.31 | 0.50 | 0.25 | 0.18 | 0.53 | 0.26 | 0.57 | 0.91 | 0.28 | 0.19 | 0.06 |
| | dV2 | 0.47 | 0.44 | 0.96 | 0.53 | 2.00 | 0.52 | 0.40 | 0.58 | 0.76 | 2.60 | 0.68 | 0.90 | 0.49 | 0.50 | 0.40 | 0.56 | 0.62 | 0.55 | 0.65 | 0.42 |
| | uV2 | 0.48 | 0.45 | 1.00 | 0.41 | 0.76 | 0.40 | 0.51 | 0.57 | 1.06 | 0.76 | 0.59 | 0.49 | 0.74 | 0.52 | 0.49 | 0.80 | 0.81 | 0.51 | 0.30 | 0.44 |
| | dV3 | 0.32 | 0.50 | 0.87 | 0.37 | 0.54 | 0.55 | 0.29 | 0.64 | 0.95 | 1.27 | 0.61 | 2.06 | 0.29 | 0.62 | 0.50 | 1.00 | 0.66 | 0.26 | 0.40 | 0.47 |
| | uV3 | 0.41 | 2.03 | 1.23 | 0.43 | 1.89 | 0.35 | 0.28 | 1.53 | 0.36 | 1.08 | 0.67 | 0.90 | 0.44 | 0.63 | 0.45 | 0.73 | 1.73 | 0.74 | 0.42 | 0.35 |
| | dV4 | 0.42 | 0.39 | 0.68 | 0.25 | 0.67 | 0.43 | 0.20 | 0.49 | 0.41 | 0.78 | 0.65 | 0.45 | 0.21 | 0.48 | 0.13 | 0.75 | 0.81 | 0.34 | 0.64 | 0.20 |
| | uV4 | 0.35 | 0.32 | 1.26 | 0.37 | 0.91 | 0.77 | 0.26 | 0.53 | 0.67 | 2.88 | 0.65 | 1.53 | 0.56 | 0.82 | 0.44 | 0.83 | 1.07 | 0.59 | 0.54 | 0.77 |
| LK-DFBA (NLR) | WT | 0.12 | 0.14 | 1.34 | 0.17 | 0.98 | 0.32 | 0.14 | 0.22 | 0.19 | 0.37 | 0.32 | 0.25 | 0.19 | 0.40 | 0.21 | 0.83 | 0.29 | 0.18 | 0.21 | 0.09 |
| | dV2 | 0.34 | 0.47 | 0.55 | 0.41 | 2.65 | 0.32 | 0.38 | 0.55 | 0.33 | 0.87 | 0.56 | 1.48 | 0.38 | 0.33 | 0.34 | 0.23 | 0.68 | 0.40 | 0.43 | 0.38 |
| | uV2 | 0.12 | 0.27 | 1.42 | 0.21 | 1.44 | 0.49 | 0.18 | 0.73 | 0.18 | 0.24 | 0.35 | 0.54 | 0.28 | 0.24 | 0.40 | 0.91 | 0.31 | 0.22 | 0.12 | 0.16 |
| | dV3 | 0.37 | 0.46 | 0.97 | 0.42 | 0.52 | 0.32 | 0.31 | 0.66 | 0.30 | 0.45 | 0.36 | 2.57 | 0.36 | 0.29 | 0.61 | 0.98 | 0.45 | 0.61 | 0.33 | 0.69 |
| | uV3 | 0.45 | 2.13 | 2.07 | 0.34 | 2.37 | 0.38 | 0.29 | 1.66 | 0.39 | 0.94 | 0.49 | 0.89 | 0.42 | 0.50 | 0.37 | 1.39 | 0.82 | 0.57 | 0.42 | 0.35 |
| | dV4 | 0.21 | 0.35 | 0.79 | 0.30 | 0.69 | 0.22 | 0.19 | 0.51 | 0.29 | 0.26 | 0.33 | 0.38 | 0.26 | 0.25 | 0.15 | 0.63 | 0.44 | 0.39 | 0.44 | 0.21 |
| | uV4 | 0.39 | 0.23 | 2.31 | 0.35 | 1.31 | 0.77 | 0.24 | 0.39 | 0.61 | 1.03 | 0.52 | 0.87 | 0.51 | 0.68 | 0.40 | 1.42 | 0.90 | 0.61 | 0.64 | 0.56 |
| LK-DFBA (DR) | WT | 0.69 | 0.40 | 0.35 | 0.37 | 0.37 | 0.20 | 0.45 | 0.49 | 0.23 | 1.45 | 0.93 | 0.21 | 0.37 | 0.60 | 0.26 | 0.82 | 0.46 | 0.15 | 0.26 | 0.19 |
| | dV2 | 0.49 | 0.58 | 0.32 | 0.47 | 0.63 | 0.38 | 0.49 | 0.67 | 0.60 | 2.74 | 0.63 | 0.59 | 0.40 | 0.58 | 0.39 | 0.62 | 0.69 | 0.45 | 0.64 | 0.37 |
| | uV2 | 0.89 | 0.64 | 0.70 | 0.63 | 0.25 | 0.25 | 0.57 | 0.75 | 0.82 | 1.00 | 0.84 | 0.59 | 0.74 | 0.59 | 0.49 | 1.15 | 1.19 | 0.48 | 0.62 | 0.45 |
| | dV3 | 0.66 | 0.63 | 0.37 | 0.43 | 0.32 | 0.37 | 0.24 | 0.71 | 1.07 | 1.77 | 0.86 | 0.35 | 0.47 | 0.66 | 0.57 | 1.47 | 0.65 | 0.44 | 0.39 | 1.01 |
| | uV3 | 0.89 | 0.76 | 0.39 | 0.41 | 0.57 | 0.23 | 0.33 | 0.83 | 0.66 | 0.96 | 0.66 | 1.18 | 0.67 | 0.64 | 0.60 | 0.54 | 1.43 | 0.18 | 0.45 | 0.35 |
| | dV4 | 0.37 | 0.80 | 0.23 | 0.34 | 0.24 | 0.12 | 0.11 | 0.36 | 0.27 | 0.75 | 0.68 | 0.53 | 0.13 | 0.49 | 0.14 | 0.37 | 0.48 | 0.21 | 0.38 | 0.12 |
| | uV4 | 0.78 | 0.51 | 0.47 | 0.69 | 0.47 | 0.47 | 0.53 | 0.79 | 0.97 | 4.49 | 1.54 | 0.25 | 0.74 | 1.04 | 0.40 | 1.42 | 1.22 | 1.85 | 0.55 | 1.71 |
| LK-DFBA (HP) | WT | 0.63 | 0.97 | 1.45 | 0.53 | 0.85 | 0.85 | 0.32 | 0.80 | 1.07 | 1.42 | 0.94 | 0.62 | 0.99 | 0.59 | 0.16 | 1.54 | 0.83 | 0.49 | 0.30 | 0.65 |
| | dV2 | 0.61 | 2.00 | 1.06 | 0.48 | 1.26 | 0.97 | 0.17 | 1.74 | 0.77 | 2.62 | 0.66 | 0.65 | 0.55 | 0.45 | 0.28 | 1.26 | 1.01 | 0.49 | 0.60 | 0.57 |
| | uV2 | 0.85 | 0.96 | 2.08 | 0.61 | 1.52 | 0.82 | 0.67 | 0.97 | 1.68 | 0.99 | 0.91 | 0.75 | 1.53 | 0.57 | 0.44 | 1.87 | 1.54 | 0.67 | 0.64 | 0.76 |
| | dV3 | 0.51 | 0.40 | 1.38 | 0.45 | 0.46 | 0.68 | 0.31 | 0.46 | 1.31 | 1.62 | 0.69 | 1.31 | 0.78 | 0.57 | 0.53 | 1.00 | 0.91 | 0.52 | 0.43 | 0.77 |
| | uV3 | 1.27 | 7.31 | 1.34 | 0.80 | 1.88 | 0.78 | 0.43 | 16.7 | 0.80 | 0.95 | 1.68 | 1.03 | 0.89 | 0.71 | 0.43 | 2.66 | 1.45 | 0.80 | 0.48 | 0.64 |
| | dV4 | 0.51 | 0.93 | 0.83 | 0.39 | 1.30 | 0.47 | 0.23 | 0.40 | 0.62 | 0.74 | 0.60 | 0.5 | 0.52 | 0.49 | 0.15 | 1.89 | 0.59 | 0.43 | 0.49 | 0.34 |
| | uV4 | 0.72 | 1.06 | 2.41 | 0.64 | 0.68 | 1.70 | 0.59 | 1.68 | 1.94 | 3.60 | 1.28 | 1.14 | 1.89 | 1.11 | 0.25 | 1.56 | 1.91 | 1.32 | 0.70 | 1.59 |

**Figure S3: LK-DFBA performance on noisy synthetic model data, nT = 50, CoV = 0.15** Each constraint approach was used to fit parameters to noisy (nT = 50, CoV = 0.15) wild-type (WT) data and then used to simulate the WT system and the system with *in silico* genetic perturbations with fluxes $v_2$, $v_3$, or $v_4$ down- or up-regulated. Dark green boxes represent the lowest average NRMSE (N = 10) within each phenotype for each synthetic model, while dark red boxes represent the highest average NRMSE. The cells with bolded white numbers indicate the LK-DFBA approach that best fits the WT data (also highlighted in cyan). Cells with white numbers are generally consistently green, indicating that fitting to WT data is a good indicator of which approach will be optimal across perturbations.

| | | Model # | | | | | | | | | | | | | | | | | | | |
|---|---|---|---|---|---|---|---|---|---|---|---|---|---|---|---|---|---|---|---|---|---|
| | | 1 | 2 | 3 | 4 | 5 | 6 | 7 | 8 | 9 | 10 | 11 | 12 | 13 | 14 | 15 | 16 | 17 | 18 | 19 | 20 |
| LK-DFBA (LR) | WT | 0.12 | 0.14 | 0.89 | 0.14 | 0.95 | 0.40 | 0.10 | 0.32 | 0.18 | 0.98 | 0.50 | 0.20 | 0.17 | 0.55 | 0.18 | 0.54 | 0.92 | 0.15 | 0.12 | 0.06 |
| | dV2 | 0.46 | 0.38 | 0.62 | 0.47 | 1.51 | 0.49 | 0.43 | 0.61 | 0.63 | 1.96 | 0.67 | 0.95 | 0.44 | 0.52 | 0.39 | 0.57 | 0.66 | 0.46 | 0.50 | 0.50 |
| | uV2 | 0.47 | 0.24 | 1.37 | 0.40 | 0.85 | 0.32 | 0.53 | 0.47 | 0.84 | 0.61 | 0.56 | 0.47 | 0.71 | 0.52 | 0.48 | 0.58 | 0.87 | 0.48 | 0.35 | 0.32 |
| | dV3 | 0.19 | 0.49 | 1.07 | 0.40 | 0.52 | 0.56 | 0.31 | 0.62 | 0.60 | 0.96 | 0.60 | 2.05 | 0.31 | 0.65 | 0.55 | 0.83 | 0.53 | 0.29 | 0.31 | 0.65 |
| | uV3 | 0.48 | 2.24 | 0.94 | 0.43 | 1.81 | 0.32 | 0.26 | 1.58 | 0.37 | 1.02 | 0.60 | 0.87 | 0.47 | 0.63 | 0.38 | 0.80 | 1.88 | 0.68 | 0.30 | 0.36 |
| | dV4 | 0.54 | 0.30 | 0.70 | 0.35 | 0.58 | 0.45 | 0.27 | 0.52 | 0.19 | 0.64 | 0.55 | 0.40 | 0.23 | 0.49 | 0.10 | 0.83 | 0.70 | 0.29 | 0.61 | 0.21 |
| | uV4 | 0.36 | 0.25 | 1.57 | 0.40 | 1.22 | 0.75 | 0.29 | 0.58 | 0.59 | 2.36 | 0.65 | 1.65 | 0.56 | 0.79 | 0.39 | 0.69 | 0.91 | 0.71 | 0.51 | 0.82 |
| LK-DFBA (NLR) | WT | 0.12 | 0.13 | 0.94 | 0.11 | 0.96 | 0.31 | 0.17 | 0.22 | 0.16 | 0.32 | 0.30 | 0.19 | 0.21 | 0.44 | 0.17 | 0.45 | 0.27 | 0.14 | 0.17 | 0.06 |
| | dV2 | 0.32 | 0.47 | 0.41 | 0.38 | 2.43 | 0.35 | 0.41 | 0.55 | 0.26 | 0.81 | 0.52 | 1.82 | 0.35 | 0.31 | 0.31 | 0.28 | 0.63 | 0.34 | 0.38 | 0.36 |
| | uV2 | 0.17 | 0.18 | 1.41 | 0.38 | 1.06 | 0.34 | 0.20 | 0.69 | 0.18 | 0.24 | 0.34 | 0.52 | 0.35 | 0.29 | 0.27 | 0.78 | 0.32 | 0.22 | 0.19 | 0.12 |
| | dV3 | 0.30 | 0.46 | 0.96 | 0.30 | 0.51 | 0.32 | 0.37 | 0.65 | 0.33 | 0.46 | 0.37 | 2.70 | 0.34 | 0.58 | 0.73 | 0.79 | 0.44 | 0.53 | 0.20 | 0.74 |
| | uV3 | 0.48 | 2.26 | 1.13 | 0.39 | 2.25 | 0.38 | 0.26 | 1.75 | 0.40 | 0.98 | 0.48 | 0.88 | 0.53 | 0.52 | 0.37 | 0.94 | 0.83 | 0.64 | 0.33 | 0.36 |
| | dV4 | 0.22 | 0.30 | 0.56 | 0.32 | 0.53 | 0.24 | 0.26 | 0.54 | 0.31 | 0.24 | 0.33 | 0.39 | 0.26 | 0.23 | 0.11 | 0.55 | 0.44 | 0.43 | 0.41 | 0.24 |
| | uV4 | 0.43 | 0.24 | 1.73 | 0.37 | 1.55 | 0.74 | 0.35 | 0.47 | 0.57 | 0.90 | 0.52 | 0.86 | 0.58 | 0.72 | 0.37 | 0.84 | 0.91 | 0.71 | 0.58 | 0.80 |
| LK-DFBA (DR) | WT | 0.70 | 0.39 | 0.55 | 0.30 | 0.28 | 0.22 | 0.38 | 0.72 | 0.14 | 1.67 | 0.90 | 0.18 | 0.42 | 0.66 | 0.24 | 0.85 | 0.22 | 0.20 | 0.40 | 0.19 |
| | dV2 | 0.46 | 0.47 | 0.51 | 0.42 | 0.44 | 0.37 | 0.50 | 0.68 | 0.46 | 2.75 | 0.62 | 0.33 | 0.34 | 0.63 | 0.30 | 0.75 | 0.72 | 0.39 | 0.47 | 0.31 |
| | uV2 | 0.80 | 0.48 | 0.75 | 0.69 | 0.11 | 0.24 | 0.51 | 0.78 | 0.69 | 1.38 | 0.82 | 0.33 | 0.76 | 0.64 | 0.48 | 1.10 | 1.04 | 0.48 | 0.55 | 0.45 |
| | dV3 | 0.70 | 0.60 | 0.57 | 0.60 | 0.30 | 0.35 | 0.33 | 0.76 | 1.33 | 1.98 | 0.90 | 3.22 | 0.56 | 0.68 | 0.53 | 1.48 | 0.39 | 0.51 | 0.44 | 1.17 |
| | uV3 | 0.86 | 0.75 | 0.57 | 0.32 | 0.43 | 0.23 | 0.45 | 0.81 | 0.75 | 1.14 | 0.68 | 1.68 | 0.79 | 0.71 | 0.62 | 0.65 | 0.98 | 0.28 | 0.46 | 0.34 |
| | dV4 | 0.52 | 0.77 | 0.50 | 0.31 | 0.18 | 0.11 | 0.18 | 0.41 | 0.47 | 0.93 | 0.67 | 0.87 | 0.17 | 0.52 | 0.12 | 0.56 | 0.45 | 0.15 | 0.17 | 0.11 |
| | uV4 | 0.72 | 0.52 | 0.66 | 0.78 | 0.48 | 0.49 | 0.68 | 1.17 | 0.65 | 3.81 | 1.46 | 0.4 | 0.56 | 1.10 | 0.42 | 1.44 | 0.81 | 2.18 | 0.67 | 1.86 |
| LK-DFBA (HP) | WT | 0.44 | 1.49 | 1.97 | 0.71 | 1.81 | 0.88 | 0.44 | 1.24 | 2.04 | 2.18 | 0.91 | 1.88 | 1.38 | 0.89 | 0.25 | 1.10 | 1.62 | 1.01 | 0.38 | 0.83 |
| | dV2 | 0.58 | 2.54 | 1.51 | 0.60 | 0.83 | 1.05 | 0.32 | 1.77 | 1.26 | 3.66 | 0.72 | 3.04 | 1.04 | 0.90 | 0.22 | 1.33 | 1.49 | 0.98 | 0.49 | 0.89 |
| | uV2 | 0.52 | 1.60 | 2.44 | 0.77 | 1.67 | 1.41 | 0.75 | 1.05 | 3.17 | 1.75 | 1.00 | 1.40 | 1.60 | 0.81 | 0.34 | 1.40 | 1.59 | 1.15 | 0.51 | 0.82 |
| | dV3 | 0.69 | 0.48 | 2.31 | 0.58 | 0.95 | 0.73 | 0.72 | 0.71 | 2.36 | 1.80 | 0.73 | 4.84 | 1.22 | 0.75 | 0.31 | 0.85 | 1.58 | 1.09 | 0.58 | 1.17 |
| | uV3 | 0.33 | 9.52 | 1.57 | 1.07 | 3.85 | 1.05 | 0.53 | 14.1 | 1.22 | 2.00 | 1.62 | 1.28 | 1.27 | 0.87 | 0.56 | 1.87 | 2.35 | 0.93 | 0.73 | 0.65 |
| | dV4 | 0.26 | 1.71 | 1.05 | 0.63 | 1.55 | 0.60 | 0.23 | 0.93 | 1.11 | 1.20 | 0.78 | 1.3 | 0.75 | 0.61 | 0.21 | 1.38 | 0.88 | 0.62 | 0.33 | 0.46 |
| | uV4 | 0.76 | 1.20 | 3.46 | 0.79 | 1.87 | 1.58 | 0.88 | 1.40 | 3.66 | 4.45 | 1.11 | 4.0 | 2.49 | 1.50 | 0.38 | 1.70 | 4.14 | 3.08 | 0.61 | 2.07 |

**Figure S4: LK-DFBA performance on noisy synthetic model data, nT = 15, CoV = 0.05** Each constraint approach was used to fit parameters to noisy (nT = 15, CoV = 0.05) wild-type (WT) data and then used to simulate the WT system and the system with *in silico* genetic perturbations with fluxes $v_2$, $v_3$, or $v_4$ down- or up-regulated. Dark green boxes represent the lowest average NRMSE (N = 10) within each phenotype for each synthetic model, while dark red boxes represent the highest average NRMSE. The cells with bolded white numbers indicate the LK-DFBA approach that best fits the WT data (also highlighted in cyan). Cells with white numbers are generally consistently green, indicating that fitting to WT data is a good indicator of which approach will be optimal across perturbations.

| | | Model # | | | | | | | | | | | | | | | | | | | |
|---|---|---|---|---|---|---|---|---|---|---|---|---|---|---|---|---|---|---|---|---|---|
| | | 1 | 2 | 3 | 4 | 5 | 6 | 7 | 8 | 9 | 10 | 11 | 12 | 13 | 14 | 15 | 16 | 17 | 18 | 19 | 20 |
| LK-DFBA (LR) | WT | 0.11 | 0.17 | 0.86 | 0.27 | 0.55 | 0.36 | 0.11 | 0.31 | 0.24 | 0.76 | 0.50 | 0.20 | 0.18 | 0.54 | 0.29 | 0.57 | 0.76 | 0.30 | 0.16 | 0.08 |
| | dV2 | 0.44 | 0.47 | 0.83 | 0.50 | 1.25 | 0.48 | 0.44 | 0.60 | 0.63 | 1.40 | 0.66 | 0.92 | 0.44 | 0.52 | 0.43 | 0.56 | 0.67 | 0.50 | 0.52 | 0.54 |
| | uV2 | 0.46 | 0.23 | 0.90 | 0.41 | 0.61 | 0.39 | 0.52 | 0.51 | 0.96 | 0.58 | 0.58 | 0.50 | 0.68 | 0.52 | 0.49 | 0.66 | 0.89 | 0.52 | 0.40 | 0.34 |
| | dV3 | 0.20 | 0.51 | 0.82 | 0.47 | 0.34 | 0.49 | 0.32 | 0.62 | 0.61 | 0.82 | 0.59 | 1.92 | 0.30 | 0.64 | 0.61 | 0.87 | 0.63 | 0.36 | 0.41 | 0.59 |
| | uV3 | 0.47 | 2.14 | 1.21 | 0.47 | 1.34 | 0.31 | 0.26 | 1.63 | 0.43 | 1.00 | 0.69 | 0.85 | 0.46 | 0.62 | 0.45 | 0.83 | 1.47 | 0.80 | 0.35 | 0.35 |
| | dV4 | 0.48 | 0.39 | 0.74 | 0.43 | 0.60 | 0.44 | 0.24 | 0.49 | 0.43 | 0.63 | 0.60 | 0.40 | 0.26 | 0.50 | 0.16 | 0.77 | 0.75 | 0.46 | 0.52 | 0.22 |
| | uV4 | 0.35 | 0.25 | 1.19 | 0.46 | 0.62 | 0.72 | 0.28 | 0.49 | 0.61 | 1.62 | 0.61 | 1.24 | 0.58 | 0.83 | 0.46 | 0.74 | 1.11 | 0.70 | 0.55 | 0.91 |
| LK-DFBA (NLR) | WT | 0.18 | 0.17 | 1.22 | 0.25 | 0.64 | 0.35 | 0.18 | 0.21 | 0.20 | 0.73 | 0.31 | 0.41 | 0.24 | 0.43 | 0.21 | 0.66 | 0.52 | 0.21 | 0.18 | 0.14 |
| | dV2 | 0.36 | 0.44 | 0.78 | 0.42 | 2.48 | 0.35 | 0.44 | 0.48 | 0.29 | 1.39 | 0.50 | 1.69 | 0.37 | 0.36 | 0.32 | 0.32 | 0.84 | 0.38 | 0.36 | 0.39 |
| | uV2 | 0.23 | 0.17 | 1.51 | 0.30 | 1.08 | 0.55 | 0.28 | 0.44 | 0.18 | 0.43 | 0.38 | 0.54 | 0.33 | 0.32 | 0.32 | 0.85 | 1.15 | 0.33 | 0.27 | 0.14 |
| | dV3 | 0.30 | 0.50 | 0.98 | 0.34 | 0.38 | 0.35 | 0.36 | 0.55 | 0.40 | 0.59 | 0.34 | 2.51 | 0.35 | 0.47 | 0.64 | 0.80 | 0.57 | 0.53 | 0.33 | 0.66 |
| | uV3 | 0.44 | 2.10 | 1.95 | 0.41 | 1.48 | 0.38 | 0.26 | 2.29 | 0.38 | 1.71 | 0.52 | 1.09 | 0.53 | 0.50 | 0.36 | 1.26 | 1.11 | 0.57 | 0.44 | 0.36 |
| | dV4 | 0.37 | 0.34 | 0.85 | 0.28 | 0.54 | 0.27 | 0.23 | 0.43 | 0.29 | 0.46 | 0.31 | 0.41 | 0.30 | 0.26 | 0.16 | 0.67 | 0.53 | 0.40 | 0.53 | 0.24 |
| | uV4 | 0.36 | 0.24 | 2.18 | 0.45 | 0.84 | 0.83 | 0.34 | 0.38 | 0.54 | 1.77 | 0.48 | 0.95 | 0.64 | 0.70 | 0.38 | 1.11 | 1.05 | 0.67 | 0.64 | 0.73 |
| LK-DFBA (DR) | WT | 0.64 | 0.39 | 0.59 | 0.48 | 0.34 | 0.21 | 0.40 | 0.58 | 0.29 | 1.38 | 0.87 | 0.21 | 0.42 | 0.62 | 0.29 | 0.77 | 0.52 | 0.31 | 0.42 | 0.24 |
| | dV2 | 0.43 | 0.46 | 0.57 | 0.47 | 0.49 | 0.34 | 0.49 | 0.64 | 0.51 | 1.99 | 0.62 | 0.35 | 0.39 | 0.60 | 0.38 | 0.67 | 0.64 | 0.42 | 0.51 | 0.34 |
| | uV2 | 0.82 | 0.47 | 0.77 | 0.70 | 0.25 | 0.26 | 0.53 | 0.76 | 0.89 | 1.19 | 0.81 | 0.40 | 0.74 | 0.61 | 0.49 | 1.04 | 0.91 | 0.48 | 0.63 | 0.46 |
| | dV3 | 0.62 | 0.60 | 0.60 | 0.71 | 0.44 | 0.34 | 0.31 | 0.73 | 1.04 | 1.62 | 0.91 | 1.80 | 0.60 | 0.70 | 0.58 | 1.39 | 0.60 | 0.61 | 0.46 | 1.12 |
| | uV3 | 0.82 | 0.76 | 0.62 | 0.50 | 0.55 | 0.23 | 0.39 | 0.84 | 0.78 | 0.92 | 0.67 | 1.60 | 0.82 | 0.71 | 0.63 | 0.57 | 1.30 | 0.42 | 0.49 | 0.36 |
| | dV4 | 0.40 | 0.76 | 0.54 | 0.38 | 0.25 | 0.12 | 0.14 | 0.32 | 0.46 | 0.87 | 0.66 | 0.70 | 0.21 | 0.51 | 0.18 | 0.47 | 0.53 | 0.23 | 0.23 | 0.15 |
| | uV4 | 0.74 | 0.49 | 0.69 | 0.82 | 0.62 | 0.46 | 0.63 | 0.84 | 0.66 | 2.73 | 1.40 | 0.37 | 0.93 | 1.12 | 0.44 | 1.37 | 1.15 | 2.22 | 0.65 | 1.79 |
| LK-DFBA (HP) | WT | 0.72 | 1.54 | 1.73 | 0.71 | 2.33 | 1.33 | 0.63 | 1.30 | 1.93 | 1.51 | 0.91 | 1.96 | 1.54 | 0.80 | 0.51 | 1.25 | 0.74 | 1.06 | 1.05 | 0.63 |
| | dV2 | 0.86 | 2.54 | 1.32 | 0.88 | 1.68 | 1.21 | 0.57 | 2.11 | 1.27 | 2.35 | 0.86 | 2.11 | 1.14 | 0.76 | 0.44 | 1.19 | 1.25 | 1.10 | 0.87 | 0.64 |
| | uV2 | 0.73 | 1.35 | 2.21 | 0.66 | 1.80 | 1.54 | 0.86 | 1.05 | 2.86 | 1.46 | 0.84 | 1.92 | 1.82 | 0.76 | 0.61 | 1.64 | 1.00 | 1.16 | 1.04 | 0.68 |
| | dV3 | 0.71 | 0.69 | 1.98 | 0.60 | 1.46 | 1.09 | 0.68 | 0.77 | 2.42 | 1.76 | 0.77 | 4.89 | 1.25 | 0.75 | 0.46 | 0.84 | 0.95 | 1.35 | 1.02 | 1.04 |
| | uV3 | 1.28 | 7.93 | 1.47 | 1.15 | 4.44 | 1.54 | 0.68 | 16.1 | 1.17 | 1.13 | 1.53 | 1.08 | 1.52 | 0.83 | 0.66 | 2.26 | 1.19 | 1.04 | 1.46 | 0.70 |
| | dV4 | 0.83 | 1.48 | 1.02 | 0.80 | 2.29 | 0.81 | 0.47 | 1.31 | 1.08 | 0.96 | 0.72 | 0.8 | 0.84 | 0.62 | 0.55 | 1.86 | 0.77 | 0.68 | 0.80 | 0.62 |
| | uV4 | 0.77 | 1.49 | 2.96 | 0.74 | 2.69 | 2.66 | 0.92 | 1.45 | 3.38 | 2.71 | 1.19 | 6.58 | 2.79 | 1.33 | 0.56 | 1.55 | 1.64 | 2.80 | 2.08 | 1.61 |

**Figure S5: LK-DFBA performance on noisy synthetic model data, nT = 15, CoV = 0.15** Each constraint approach was used to fit parameters to noisy (nT = 15, CoV = 0.15) wild-type (WT) data and then used to simulate the WT system and the system with *in silico* genetic perturbations with fluxes $v_2$, $v_3$, or $v_4$ down- or up-regulated. Dark green boxes represent the lowest average NRMSE (N = 10) within each phenotype for each synthetic model, while dark red boxes represent the highest average NRMSE. The cells with bolded white numbers indicate the LK-DFBA approach that best fits the WT data (also highlighted in cyan). Cells with white numbers are generally consistently green, indicating that fitting to WT data is a good indicator of which approach will be optimal across perturbations.

| | | Model # | | | | | | | | | | | | | | | | | | | |
|---|---|---|---|---|---|---|---|---|---|---|---|---|---|---|---|---|---|---|---|---|---|
| | | 1 | 2 | 3 | 4 | 5 | 6 | 7 | 8 | 9 | 10 | 11 | 12 | 13 | 14 | 15 | 16 | 17 | 18 | 19 | 20 |
| LK-DFBA (LR) | WT | 0.10 | 0.18 | 0.74 | 0.23 | 0.63 | 0.40 | 0.12 | 0.34 | 0.29 | 0.79 | 0.51 | 0.30 | 0.18 | 0.54 | 0.35 | 0.62 | 1.10 | 0.32 | 0.17 | 0.08 |
| | dV2 | 0.43 | 0.51 | 0.65 | 0.49 | 1.14 | 0.50 | 0.41 | 0.62 | 0.62 | 1.40 | 0.67 | 0.84 | 0.46 | 0.52 | 0.44 | 0.56 | 0.65 | 0.56 | 0.55 | 0.50 |
| | uV2 | 0.47 | 0.32 | 0.97 | 0.41 | 0.70 | 0.31 | 0.49 | 0.47 | 0.99 | 0.61 | 0.58 | 0.56 | 0.72 | 0.52 | 0.53 | 1.40 | 1.02 | 0.51 | 0.49 | 0.32 |
| | dV3 | 0.19 | 0.50 | 0.88 | 0.48 | 0.37 | 0.56 | 0.34 | 0.65 | 0.72 | 0.90 | 0.63 | 2.21 | 0.32 | 0.63 | 0.55 | 0.95 | 0.76 | 0.38 | 0.34 | 0.62 |
| | uV3 | 0.45 | 2.20 | 1.03 | 0.47 | 1.31 | 0.34 | 0.27 | 1.58 | 0.43 | 0.83 | 0.66 | 0.84 | 0.45 | 0.63 | 0.48 | 0.99 | 2.11 | 0.79 | 0.45 | 0.36 |
| | dV4 | 0.44 | 0.49 | 0.71 | 0.38 | 0.56 | 0.47 | 0.23 | 0.52 | 0.35 | 0.67 | 0.58 | 0.47 | 0.24 | 0.49 | 0.26 | 0.85 | 0.83 | 0.46 | 0.53 | 0.22 |
| | uV4 | 0.33 | 0.26 | 1.19 | 0.46 | 0.55 | 0.75 | 0.28 | 0.51 | 0.64 | 1.84 | 0.64 | 1.55 | 0.56 | 0.81 | 0.45 | 0.81 | 1.50 | 0.72 | 0.56 | 0.77 |
| LK-DFBA (NLR) | WT | 0.19 | 0.16 | 1.39 | 0.21 | 0.68 | 0.34 | 0.18 | 0.23 | 0.23 | 0.56 | 0.37 | 0.33 | 0.21 | 0.41 | 0.25 | 0.77 | 0.37 | 0.26 | 0.19 | 0.13 |
| | dV2 | 0.37 | 0.48 | 0.47 | 0.42 | 2.29 | 0.36 | 0.47 | 0.55 | 0.31 | 1.14 | 0.53 | 1.53 | 0.39 | 0.38 | 0.33 | 0.31 | 0.75 | 0.43 | 0.38 | 0.43 |
| | uV2 | 0.30 | 0.19 | 1.61 | 0.29 | 1.03 | 0.38 | 0.29 | 0.59 | 0.25 | 0.42 | 0.38 | 0.52 | 0.38 | 0.29 | 0.39 | 1.47 | 0.60 | 0.32 | 0.20 | 0.24 |
| | dV3 | 0.37 | 0.46 | 0.99 | 0.39 | 0.37 | 0.35 | 0.44 | 0.59 | 0.39 | 0.66 | 0.38 | 2.84 | 0.37 | 0.46 | 0.58 | 0.98 | 0.43 | 0.47 | 0.30 | 0.81 |
| | uV3 | 0.44 | 2.04 | 2.38 | 0.41 | 1.49 | 0.38 | 0.28 | 2.33 | 0.43 | 1.45 | 0.50 | 0.95 | 0.47 | 0.51 | 0.39 | 1.37 | 0.77 | 0.60 | 0.48 | 0.34 |
| | dV4 | 0.32 | 0.39 | 0.82 | 0.30 | 0.50 | 0.26 | 0.26 | 0.50 | 0.31 | 0.35 | 0.30 | 0.39 | 0.28 | 0.26 | 0.21 | 1.00 | 0.40 | 0.37 | 0.40 | 0.25 |
| | uV4 | 0.41 | 0.22 | 2.41 | 0.44 | 0.80 | 0.79 | 0.34 | 0.44 | 0.55 | 1.34 | 0.61 | 1.31 | 0.57 | 0.68 | 0.40 | 1.27 | 0.97 | 0.71 | 0.63 | 0.62 |
| LK-DFBA (DR) | WT | 0.55 | 0.39 | 0.43 | 0.53 | 0.88 | 0.22 | 0.43 | 0.56 | 0.37 | 1.70 | 0.90 | 0.38 | 0.36 | 0.64 | 0.30 | 0.69 | 0.77 | 0.36 | 0.42 | 0.21 |
| | dV2 | 0.41 | 0.51 | 0.38 | 0.50 | 0.58 | 0.35 | 0.49 | 0.68 | 0.53 | 2.77 | 0.64 | 0.72 | 0.40 | 0.64 | 0.40 | 0.57 | 0.76 | 0.48 | 0.53 | 0.35 |
| | uV2 | 0.80 | 0.55 | 0.71 | 0.67 | 0.90 | 0.27 | 0.54 | 0.78 | 0.99 | 1.33 | 0.80 | 0.64 | 0.71 | 0.63 | 0.49 | 1.06 | 1.22 | 0.49 | 0.68 | 0.45 |
| | dV3 | 0.52 | 0.59 | 0.45 | 0.64 | 0.69 | 0.36 | 0.31 | 0.77 | 1.11 | 1.61 | 0.87 | 2.64 | 0.54 | 0.68 | 0.59 | 1.38 | 0.52 | 0.64 | 0.47 | 1.10 |
| | uV3 | 0.73 | 0.75 | 0.47 | 0.55 | 1.63 | 0.26 | 0.42 | 0.90 | 0.79 | 0.92 | 0.70 | 1.87 | 0.68 | 0.71 | 0.66 | 0.52 | 1.41 | 0.41 | 0.50 | 0.36 |
| | dV4 | 0.33 | 0.79 | 0.36 | 0.38 | 0.53 | 0.13 | 0.16 | 0.37 | 0.47 | 0.96 | 0.67 | 0.7 | 0.16 | 0.51 | 0.23 | 0.35 | 0.56 | 0.27 | 0.23 | 0.15 |
| | uV4 | 0.66 | 0.49 | 0.51 | 0.86 | 1.54 | 0.44 | 0.63 | 0.86 | 0.89 | 3.60 | 1.49 | 0.80 | 0.84 | 1.04 | 0.46 | 1.30 | 1.40 | 2.20 | 0.66 | 1.76 |
| LK-DFBA (HP) | WT | 0.61 | 1.28 | 1.45 | 0.74 | 2.46 | 1.14 | 0.55 | 1.37 | 1.77 | 1.48 | 0.98 | 1.29 | 1.32 | 0.81 | 0.55 | 1.59 | 1.18 | 0.94 | 0.64 | 0.80 |
| | dV2 | 0.66 | 2.16 | 1.08 | 0.80 | 2.67 | 1.29 | 0.42 | 2.26 | 1.02 | 2.10 | 0.71 | 1.52 | 0.87 | 0.81 | 0.49 | 1.44 | 1.76 | 1.03 | 0.58 | 0.81 |
| | uV2 | 0.62 | 1.17 | 2.15 | 0.70 | 2.00 | 1.48 | 0.81 | 1.20 | 2.80 | 1.34 | 0.91 | 1.24 | 1.72 | 0.77 | 0.67 | 1.71 | 1.20 | 0.96 | 0.83 | 0.83 |
| | dV3 | 0.77 | 0.56 | 1.70 | 0.76 | 1.83 | 0.92 | 0.59 | 0.78 | 2.56 | 1.64 | 0.77 | 2.74 | 1.10 | 0.77 | 0.45 | 1.09 | 1.06 | 1.10 | 0.70 | 1.38 |
| | uV3 | 0.83 | 6.54 | 1.09 | 1.12 | 4.58 | 1.23 | 0.56 | 16.3 | 0.99 | 1.04 | 1.48 | 1.89 | 1.23 | 0.84 | 0.64 | 2.60 | 1.85 | 0.99 | 1.17 | 0.85 |
| | dV4 | 0.59 | 1.48 | 0.77 | 0.66 | 2.30 | 0.67 | 0.34 | 1.33 | 0.99 | 1.07 | 0.85 | 0.81 | 0.70 | 0.61 | 0.50 | 2.21 | 1.14 | 0.90 | 0.71 | 0.69 |
| | uV4 | 0.79 | 1.15 | 2.68 | 0.84 | 3.34 | 2.10 | 0.86 | 1.39 | 3.31 | 2.69 | 1.31 | 2.84 | 2.39 | 1.39 | 0.61 | 1.45 | 2.81 | 2.57 | 0.81 | 1.94 |

**Figure S6: LK-DFBA performance on noisy synthetic model data after smoothing.** Each constraint approach was used to fit parameters to noisy (nT = 15, CoV = 0.15) wild-type (WT) data that was smoothed with a previously described impulse function [5], and then used to simulate the WT system and the system with *in silico* genetic perturbations with fluxes $v_2$, $v_3$, or $v_4$ down- or up-regulated. Dark green boxes represent the lowest average NRMSE (N = 10) within each phenotype for each synthetic model, while dark red boxes represent the highest average NRMSE. The cells with bolded white numbers indicate the LK-DFBA approach that best fits the WT data (also highlighted in cyan). Cells with white numbers are generally consistently green, indicating that fitting to WT data is a good indicator of which approach will be optimal across perturbations.

**A.** *L. lactis*

| | | nT=50 CoV=0 | nT=50 CoV=0.05 | nT=50 CoV=0.15 | nT=15 CoV=0.05 | nT=15 CoV=0.15 |
|---|---|---|---|---|---|---|
| LK-DFBA (LR) | WT | 6.7E+04 | 2.2E+07 | 5.2E+06 | 9.0E+04 | 4.3E+04 |
| | dLDH | 9.4E+04 | 5.7E+04 | 3.0E+04 | 1.9E+03 | 1.8E+03 |
| | dPFK | 4.2E+04 | 1.8E+04 | 1.2E+04 | 2.9E+03 | 1.8E+03 |
| | uPFK | 6.2E+03 | 1.2E+07 | 1.4E+07 | 2.2E+05 | 1.5E+05 |
| | uACK | 5.8E+03 | 1.3E+07 | 4.1E+06 | 2.2E+04 | 5.6E+04 |
| | uMP | 2.8E+03 | 6.8E+06 | 2.5E+06 | 6.1E+04 | 3.8E+04 |
| LK-DFBA (NLR) | WT | 1.8E+04 | 1.8E+04 | **2.0E+04** | **9.6E+03** | **2.9E+04** |
| | dLDH | 6.5E+03 | 5.9E+02 | **2.8E+02** | **2.2E+03** | **3.3E+03** |
| | dPFK | 6.7E+03 | 2.3E+02 | **2.5E+02** | **4.6E+03** | **4.2E+03** |
| | uPFK | 1.9E+04 | 3.9E+04 | **4.2E+04** | **1.1E+04** | **6.2E+04** |
| | uACK | 1.7E+04 | 2.7E+04 | **1.8E+04** | **9.9E+03** | **2.8E+04** |
| | uMP | 1.2E+04 | 1.1E+04 | **7.7E+03** | **7.2E+03** | **1.8E+04** |
| LK-DFBA (DR) | WT | 1.9E+07 | 1.3E+08 | 9.8E+07 | 7.6E+07 | 8.2E+07 |
| | dLDH | 2.4E+05 | 9.8E+05 | 1.0E+06 | 6.1E+05 | 2.9E+05 |
| | dPFK | 1.0E+05 | 9.2E+04 | 2.2E+04 | 1.9E+04 | 6.3E+04 |
| | uPFK | 2.9E+07 | 2.6E+08 | 1.9E+08 | 2.1E+08 | 2.3E+08 |
| | uACK | 1.0E+07 | 1.5E+08 | 1.1E+08 | 7.4E+07 | 1.0E+08 |
| | uMP | 5.5E+06 | 6.8E+07 | 5.1E+07 | 3.8E+07 | 5.2E+07 |
| LK-DFBA (HP) | WT | **2.9E+01** | **4.3E+03** | 1.4E+06 | 1.3E+07 | 8.7E+06 |
| | dLDH | **2.1E+02** | **2.3E+03** | 2.6E+03 | 5.5E+05 | 8.3E+03 |
| | dPFK | **4.7E+01** | **2.4E+03** | 9.4E+02 | 2.9E+05 | 1.3E+04 |
| | uPFK | **5.7E+02** | **6.2E+06** | 1.5E+07 | 1.2E+07 | 1.7E+07 |
| | uACK | **8.6E+01** | **1.8E+06** | 2.4E+06 | 1.8E+07 | 6.7E+06 |
| | uMP | **1.2E+03** | **4.7E+03** | 1.0E+06 | 4.6E+06 | 3.7E+06 |

**B.** *E. coli*

| | | nT=50 CoV=0 | nT=50 CoV=0.05 | nT=50 CoV=0.15 | nT=15 CoV=0.05 | nT=15 CoV=0.15 |
|---|---|---|---|---|---|---|
| LK-DFBA (LR) | WT | 0.62 | 0.70 | 0.68 | 0.68 | 0.73 |
| | dPYK | 0.62 | 0.64 | 0.60 | 0.63 | 0.61 |
| | dPGI | 0.69 | 0.71 | 0.64 | 0.67 | 0.67 |
| | dGAPDH | 0.90 | 0.98 | 1.18 | 0.91 | 1.03 |
| | dPFK | 0.58 | 0.57 | 0.75 | 0.59 | 0.68 |
| | dTIS | 0.81 | 0.85 | 0.89 | 0.84 | 0.88 |
| | dRPE | 0.72 | 0.75 | 0.71 | 0.73 | 0.71 |
| | dPGM | 0.61 | 0.64 | 0.60 | 0.62 | 0.60 |
| LK-DFBA (NLR) | WT | **0.34** | **0.46** | **0.49** | **0.47** | **0.49** |
| | dPYK | **0.35** | **0.39** | **0.42** | **0.41** | **0.43** |
| | dPGI | **0.96** | **0.73** | **0.54** | **0.51** | **0.53** |
| | dGAPDH | **0.50** | **0.47** | **0.57** | **0.50** | **0.52** |
| | dPFK | **0.45** | **0.46** | **0.53** | **0.58** | **0.51** |
| | dTIS | **0.44** | **0.45** | **0.51** | **0.44** | **0.53** |
| | dRPE | **0.36** | **0.42** | **0.49** | **0.44** | **0.47** |
| | dPGM | **1.27** | **0.39** | **0.42** | **0.40** | **0.43** |
| LK-DFBA (DR) | WT | 2.10 | 0.86 | 0.82 | 0.88 | 0.84 |
| | dPYK | 2.10 | 0.89 | 0.70 | 0.89 | 0.73 |
| | dPGI | 0.79 | 0.74 | 0.81 | 0.81 | 0.82 |
| | dGAPDH | 2.18 | 1.77 | 1.67 | 1.92 | 1.67 |
| | dPFK | 1.81 | 0.84 | 0.79 | 0.70 | 0.85 |
| | dTIS | 1.98 | 1.06 | 1.04 | 1.13 | 1.16 |
| | dRPE | 1.75 | 0.96 | 0.74 | 0.89 | 0.87 |
| | dPGM | 2.10 | 0.88 | 0.70 | 0.89 | 0.73 |
| LK-DFBA (HP) | WT | 2.27 | 0.82 | 0.76 | 1.19 | 0.84 |
| | dPYK | 2.25 | 0.78 | 0.65 | 1.35 | 0.78 |
| | dPGI | 0.68 | 0.64 | 0.75 | 0.69 | 0.70 |
| | dGAPDH | 1.03 | 1.50 | 1.33 | 1.12 | 1.58 |
| | dPFK | 0.97 | 1.07 | 1.13 | 1.20 | 1.36 |
| | dTIS | 1.35 | 1.05 | 0.98 | 1.19 | 1.07 |
| | dRPE | 2.37 | 0.78 | 0.69 | 1.31 | 0.82 |
| | dPGM | 2.27 | 0.78 | 0.65 | 1.35 | 0.78 |

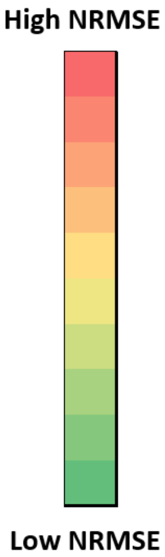


**Figure S7: LK-DFBA performance on *L. lactis* and *E. coli* models** Noiseless and noisy data performance on *L. lactis* and *E. coli* models [1, 6]. Dark green boxes represent the lowest average NRMSE (N = 10 for noisy conditions) within each phenotype for each condition, while dark red boxes represent the highest average NRMSE. The cells with bolded white numbers indicate the LK-DFBA approach that best fits the WT data (also highlighted in cyan). Cells with white numbers are generally consistently green, indicating that fitting to WT data is a good indicator of which approach will be optimal across perturbations.

| | | Model # | | | | | |
|---|---|---|---|---|---|---|---|
| | | Original | 1 | 2 | 3 | 4 | 5 |
| LK-DFBA (LR) | WT | 0.62 | 2.44 | 20.27 | **1.56** | 4.14 | 6.29 |
| | dPYK | 0.62 | 2.39 | 20.27 | **1.56** | 3.88 | 6.27 |
| | dPGI | 0.69 | 2.76 | 18.50 | **1.56** | 0.61 | 6.24 |
| | dGAPDH | 0.90 | 1.14 | 23.24 | **2.01** | 1.03 | 25.57 |
| | dPFK | 0.58 | 2.54 | 17.94 | **1.56** | 3.88 | 6.43 |
| | dTIS | 0.81 | 3.12 | 19.01 | **1.43** | 3.70 | 7.16 |
| | dRPE | 0.72 | 2.67 | 19.54 | **1.50** | 4.21 | 7.12 |
| | dPGM | 0.61 | 2.35 | 20.26 | **1.55** | 3.50 | 6.29 |
| LK-DFBA (NLR) | WT | **0.34** | **0.68** | **6.82** | 1.72 | 2.95 | 6.68 |
| | dPYK | **0.35** | **0.68** | **6.82** | 1.71 | 2.96 | 6.58 |
| | dPGI | **0.96** | **1.46** | **6.82** | 1.61 | 0.58 | 6.38 |
| | dGAPDH | **0.50** | **0.94** | **6.59** | 1.34 | 1.02 | 26.49 |
| | dPFK | **0.45** | **2.82** | **6.03** | 1.55 | 4.93 | 6.84 |
| | dTIS | **0.44** | **0.69** | **6.08** | 1.22 | 0.48 | 8.57 |
| | dRPE | **0.36** | **0.71** | **6.72** | 1.27 | 2.84 | 7.73 |
| | dPGM | **1.27** | **0.66** | **6.82** | 1.30 | 2.82 | 6.68 |
| LK-DFBA (DR) | WT | 2.10 | 1.72 | 32.25 | 1.79 | 9.86 | **1.95** |
| | dPYK | 2.10 | 1.68 | 32.25 | 1.79 | 9.86 | **1.94** |
| | dPGI | 0.79 | 6.61 | 27.57 | 1.79 | 1.34 | **2.07** |
| | dGAPDH | 2.18 | 1.96 | 41.58 | 2.03 | 3.18 | **25.40** |
| | dPFK | 1.81 | 1.14 | 18.67 | 1.79 | 6.55 | **1.87** |
| | dTIS | 1.98 | 1.98 | 29.41 | 1.81 | 6.06 | **1.76** |
| | dRPE | 1.75 | 2.39 | 31.43 | 1.79 | 9.20 | **4.25** |
| | dPGM | 2.10 | 1.66 | 32.21 | 1.50 | 9.15 | **1.95** |
| LK-DFBA (HP) | WT | 2.27 | 14.98 | 42.47 | 3.37 | **1.58** | 41.55 |
| | dPYK | 2.25 | 14.68 | 42.47 | 3.37 | **1.91** | 41.23 |
| | dPGI | 0.68 | 18.35 | 37.50 | 3.14 | **0.55** | 41.34 |
| | dGAPDH | 1.03 | 5.29 | 35.90 | 2.01 | **2.18** | 155.42 |
| | dPFK | 0.97 | 20.94 | 19.46 | 3.37 | **1.30** | 51.78 |
| | dTIS | 1.35 | 6.76 | 31.49 | 3.06 | **5.53** | 44.29 |
| | dRPE | 2.37 | 12.68 | 40.95 | 3.19 | **1.48** | 34.45 |
| | dPGM | 2.27 | 14.46 | 42.39 | 3.35 | **1.27** | 41.59 |

**Figure S8: LK-DFBA performance on different models with the same stoichiometric topology as the *E. coli* model.** Five models with the same topology as the original *E. coli* model [6] were created by randomizing the original kinetic parameters. The four LK-DFBA approaches were evaluated on noiseless data generated by these new models. Dark green boxes represent the lowest NRMSE within each phenotype for each model, while dark red boxes represent the highest NRMSE. The cells with bolded white numbers indicate the LK-DFBA approach that best fits the WT data (also highlighted in cyan). Cells with white numbers are generally consistently green, indicating that fitting to WT data is a good indicator of which approach will be optimal across perturbations.

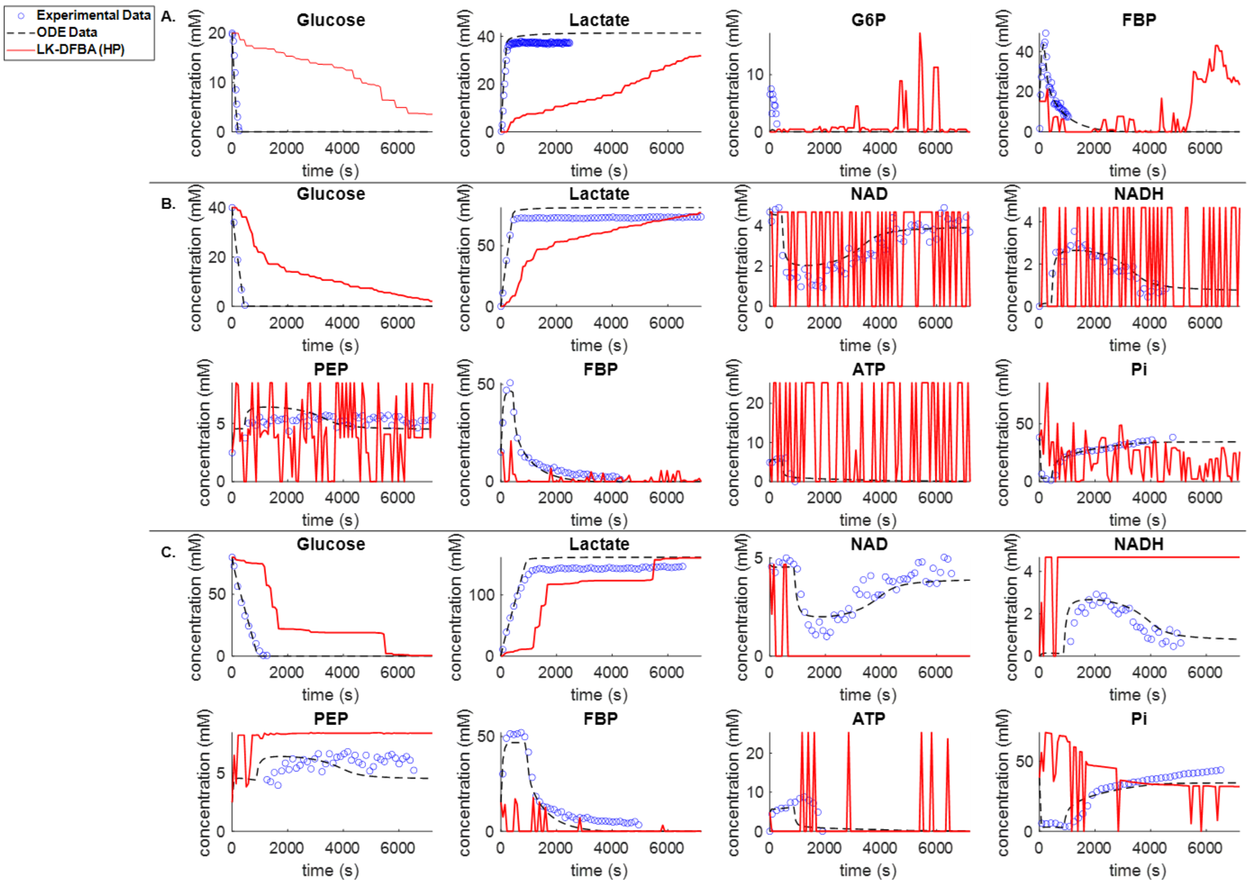


**Figure S9: Comparison of LK-DFBA metabolite concentration predictions when fitted to noiseless ODE data against noiseless ODE data and all available *L. lactis* experimental data.** Panels A, B, and C depict concentration profiles for LK-DFBA (HP) and the ODE model compared to experimental data for initial glucose concentrations of 20 mM, 40 mM, and 80 mM, respectively. Cofactor concentrations were more challenging to predict, exhibiting many spikes upwards or downwards in concentration. Cofactors were involved in several kinetics constraints and the spikes in concentrations were likely due to changes in which constraints were active in the linear program at any given point in the simulation. Nonetheless, the concentrations generally remained within an order of magnitude of the experimental data.

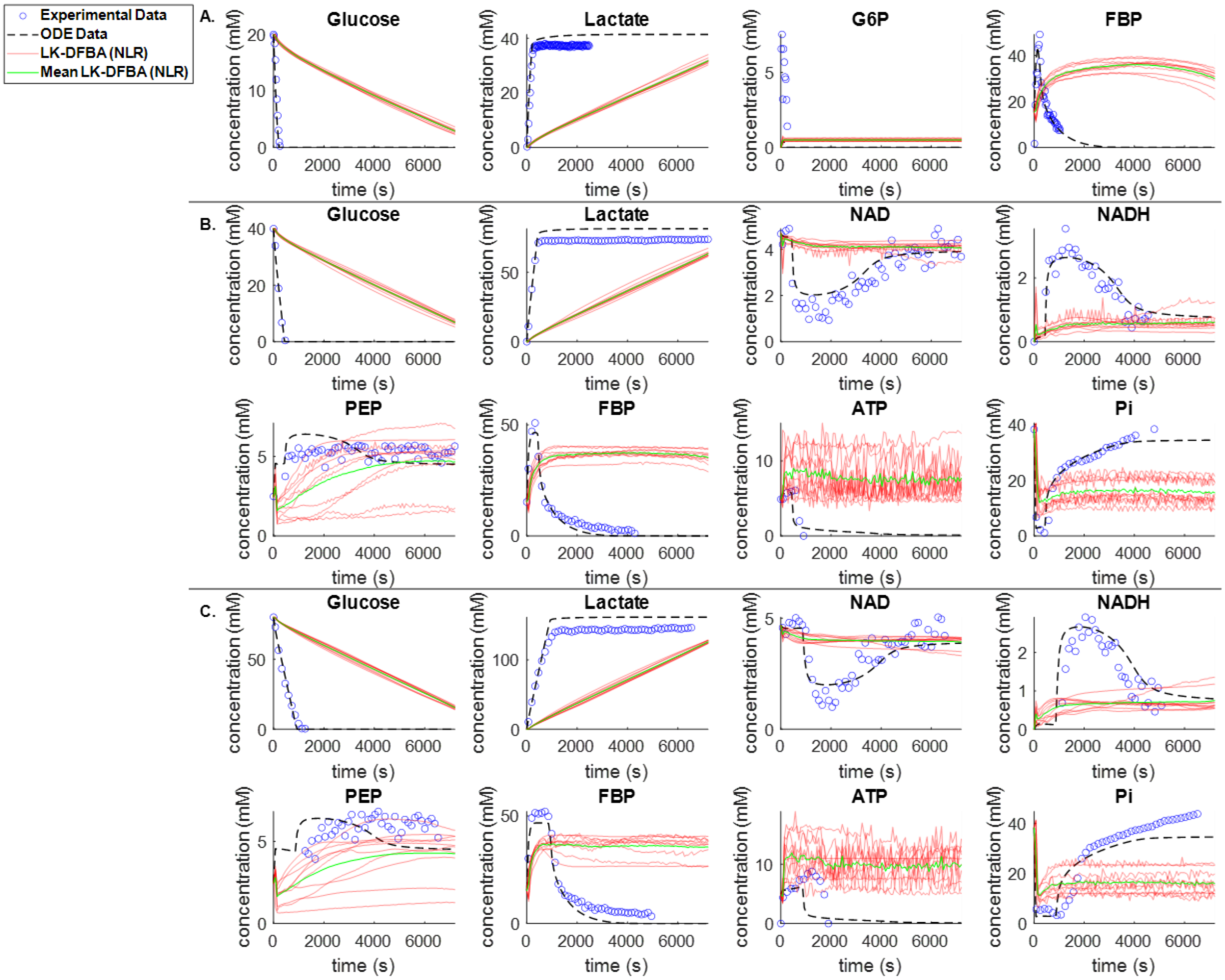


**Figure S10: Comparison of LK-DFBA metabolite concentration predictions when fitted to noisy data against ODE and all available *L. lactis* experimental data.** A., B., and C. present concentration profiles for LK-DFBA (NLR) on 10 noisy datasets (nT = 15, CoV = 0.15) and the ODE model compared to experimental data for initial glucose concentrations of 20 mM, 40 mM, and 80 mM, respectively. The mean concentration profile (solid green line) is shown with each of the concentration profiles (solid red lines) from the 10 noisy datasets.

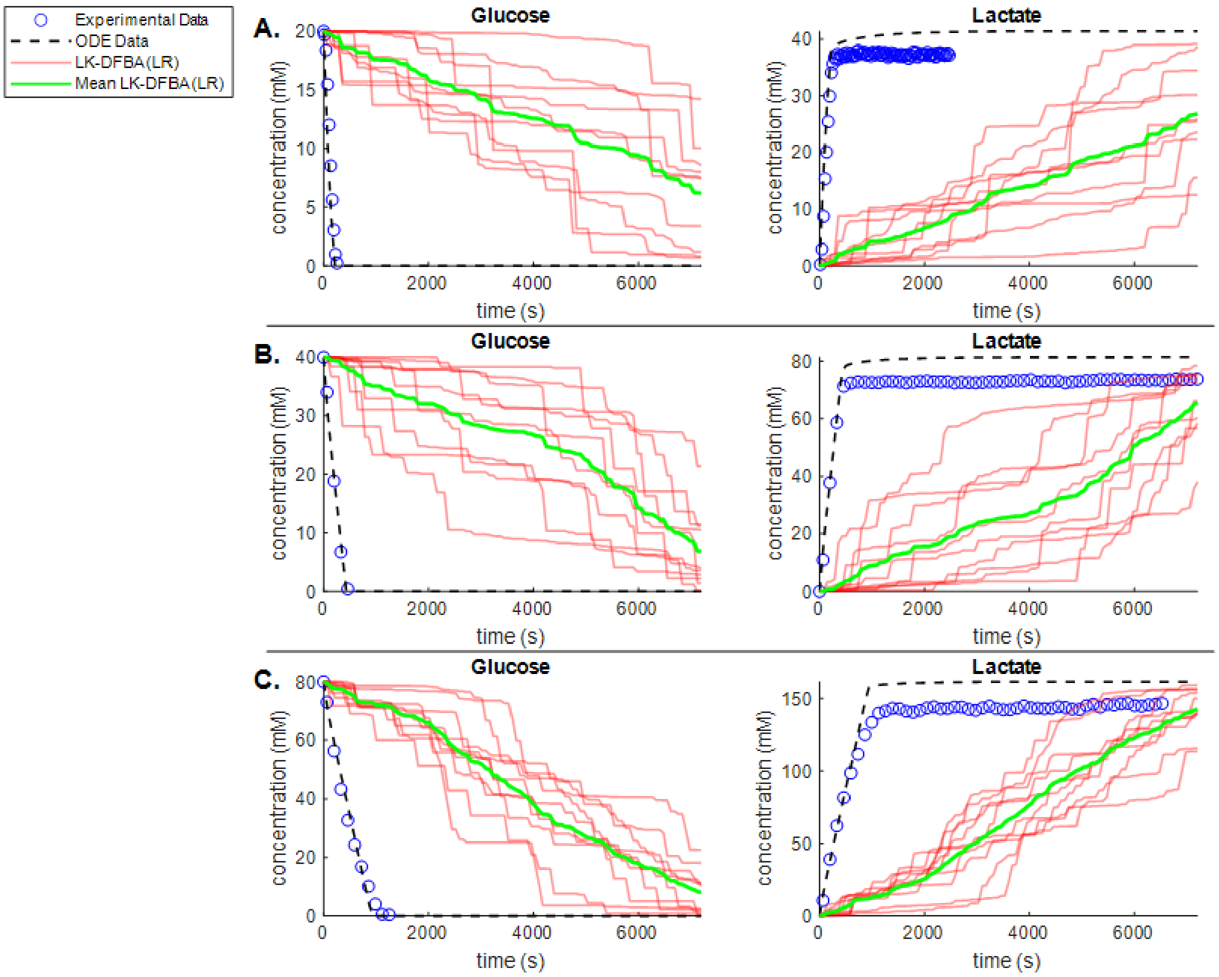


**Figure S11: Comparison of LK-DFBA (LR) metabolite concentration predictions against ODE data and *L. lactis* experimental data when fitted to noisy ODE data.** Panels A, B, and C depict concentration profiles for LK-DFBA (LR) on 10 noisy datasets (nT = 15, CoV = 0.15) and the ODE model compared to experimental data. The mean concentration profile (solid green line) is shown with each of the concentration profiles (solid red lines) from the 10 noisy datasets. While LK-DFBA (LR) is able to capture the general trends of glucose depletion and lactate accumulation, the best LK-DFBA approach on noisy data, LK-DFBA (HP), showed much more consistency in its predictions across noisy datasets for glucose and lactate (Figure 4).
